# Low-level overexpression of wild type TDP-43 causes late-onset, progressive neurodegeneration and paralysis in mice

**DOI:** 10.1101/2021.08.04.455119

**Authors:** Chunxing Yang, Tao Qiao, Jia Yu, Hongyan Wang, Yansu Guo, Johnny Salameh, Jake Metterville, Sepideh Parsi, Robert H. Brown, Huaibin Cai, Zuoshang Xu

**Affiliations:** Department of Biochemistry and Molecular Pharmacology, University of Massachusetts Medical School, Worcester, Massachusetts 01605, USA; Transgenics section, Laboratory of Neurogenetics, National Institute on Aging, National Institutes of Health, Bethesda, MD 20892, USA; Department of Neurology, University of Massachusetts Medical School, Worcester, Massachusetts 01605, USA; RNA Therapeutic Institute, University of Massachusetts Medical School, Worcester, Massachusetts 01605, USA; Neuroscience Program, University of Massachusetts Medical School, Worcester, Massachusetts 01605, USA; CoWin Biosciences, 222 Maple Avenue, Shrewsbury, MA 01545, USA; Astellas Pharma, 33 Locke Dr, Marlborough, MA 01752; Institute for Geriatrics and Rehabilitation, Beijing Geriatric Hospital, 118 Wenquan Road, Haidian District, Beijing 100095, P.R. China; Beijing Geriatric Healthcare Center, Xuanwu Hospital, Capital Medical University, No. 45 Changchun Street, Xicheng, Beijing 100053, China; Department of Neurology, American University Beirut Medical Center (AUBMC), P.O. Box 11 - 0236 / Riad El-Solh 1107 2020 Beirut, Lebanon

## Abstract

Modestly increased expression of transactive response DNA binding protein (*TDP-43*) gene have been reported in amyotrophic lateral sclerosis (ALS), frontotemporal dementia (FTD), and other neuromuscular diseases. However, whether this modest elevation triggers neurodegeneration is not known. Although high levels of TDP-43 overexpression have been modeled in mice and shown to cause early death, models with low-level overexpression that mimic the human condition have not been established. In this study, transgenic mice overexpressing wild type TDP-43 at less than 60% above the endogenous CNS levels were constructed, and their phenotypes analyzed by a variety of techniques, including biochemical, molecular, histological, behavioral techniques and electromyography. The TDP-43 transgene was expressed in neurons, astrocytes, and oligodendrocytes in the cortex and predominantly in astrocytes and oligodendrocytes in the spinal cord. The mice developed a reproducible progressive weakness ending in paralysis in mid-life. Detailed analysis showed ∼30% loss of large pyramidal neurons in the layer V motor cortex; in the spinal cord, severe demyelination was accompanied by oligodendrocyte injury, protein aggregation, astrogliosis and microgliosis, and elevation of neuroinflammation. Surprisingly, there was no loss of lower motor neurons in the lumbar spinal cord despite the complete paralysis of the hindlimbs. However, denervation was detected at the neuromuscular junction. These results demonstrate that low-level TDP-43 overexpression can cause diverse aspects of ALS, including late-onset and progressive motor dysfunction, neuroinflammation, and neurodegeneration. Our findings suggest that persistent modest elevations in TDP-43 expression can lead to ALS and other neurological disorders involving TDP-43 proteinopathy. Because of the predictable and progressive clinical paralytic phenotype, this transgenic mouse model will be useful in preclinical trial of therapeutics targeting neurological disorders associated with elevated levels of TDP-43.

## Introduction

ALS is a neurodegenerative disease that causes relentless, progressive loss of upper and lower motor neurons leading to paralysis and death. Approximately 90 percent of ALS are sporadic, whereas ∼10 percent are familial. The mechanism of motor neuron degeneration in ALS is not yet fully understood. Mutations in many genes can cause or enhance the risk of ALS. These genes function in multiple cellular mechanisms, including protein homeostasis, RNA processing, inflammation regulation, cytoskeleton dynamics, and intracellular trafficking, thus suggesting that abnormalities in diverse pathways may lead to motor neuron neurodegeneration in ALS [1, 2].

TDP-43 is an RNA binding protein whose intracellular inclusions are associated with >97% of ALS cases [3–6]. Present predominantly in the nucleus, TDP-43 regulates many RNA processes, including transcription, translation, splicing, and transport [7]. In addition, TDP-43 regulates some miRNA processing, participates in stress granule formation [8], and suppresses cryptic exon expression [9]. TDP-43 function is essential for the survival of many types of cells in mammals; ubiquitous knockout of TDP-43 results in early embryonic death in mice [10–13].

Mutations in the TDP-43 gene cause familial ALS [14–16]. These mutations increase the propensity of TDP-43 to misfold and aggregate [17] but have minimal effects on its mRNA splicing function or participation in stress granule formation [18–21]. Mutant TDP-43 proteins form insoluble intracellular aggregates in ALS patients [5]. Intriguingly, wild type TDP-43 protein also forms aggregates in sporadic ALS patients and in a large fraction of familial ALS that do not have TDP-43 mutations [3–5, 22]. This phenomenon of TDP-43 aggregation, designated as TDP-43 proteinopathy, is evident in motor neurons and oligodendrocytes [23, 24] and is thought to initiate motor neuron degeneration by both a loss of TDP-43 function and a gain of toxicity [22, 25, 26].

What drives wild type TDP-43 to proteinopathy is not understood. Several studies have suggested that modestly elevated levels of TDP-43 expression are associated with TDP-43 proteinopathy in ALS and other diseases such as FTD and inclusion body myopathy, Paget’s disease of the bone and frontotemporal dementia (IBMPFD) [27–30]. ALS-associated TDP-43 mutations stabilize the TDP-43 protein or mRNA [19, 31–33]. TDP-43 levels are increased in cells with TDP-43 mutations from ALS patients and in mutant TDP-43 knock-in transgenic animals [15, 34, 35]. Remarkably, the mutant stability is inversely correlated with the disease onset age in ALS patients; the more stable the mutant, the earlier the disease onset [19]. These studies raise the question of whether a modestly elevated level of TDP-43 suffices to trigger progressive motor neuron degeneration and ALS.

To answer this question, a mammalian model with modestly elevated levels of TDP-43 should be informative, as such a model will closely mimic the modestly elevated levels in human ALS and create an opportunity to observe chronic TDP-43 toxicity. Previous mammalian models with TDP-43 overexpression have shown a wide range of phenotypes, including both motor and non-motor symptoms [36]. In general, the high TDP-43-expressing lines die rapidly after birth. Some of these mice showed TDP-43 intracellular inclusions and mild motor neuron loss [36]. The low TDP-43 expression lines develop subtle or no motor phenotypes with only mild cellular and molecular changes during the animal’s lifespan [35–42]. These models notwithstanding, whether a modestly elevated level of TDP-43 within the range of the increases observed in human ALS can trigger neurodegeneration and ALS phenotypes remain uncertain.

In this study, we have constructed two TDP-43 transgenic lines with modest levels of TDP-43 overexpression. Both lines developed motor weakness after ∼500 days. The weakness eventually progressed to paralysis in a few months. Conversion of one line into homozygotes accelerated the disease. The homozygous mice developed paralysis during a period of 300 to 430 days. Detailed analysis showed ∼30% motor neuron loss in layer V cortex, oligodendrocyte degeneration, demyelination, and neuroinflammation, but no motor neuron loss, in the spinal cord. However, denervation at the neuromuscular junction was detected, indicating the presence of distal motor axon degeneration. Thus, this mouse model has replicated several key features of ALS, including progressive motor dysfunction, paralysis, loss of upper motor neurons, oligodendrocyte degeneration, loss of myelin and neuroinflammation. These results demonstrate that a modest elevation of TDP-43 can trigger late-onset neurodegeneration and motor dysfunction and thus may play a causative role in human ALS and other neuromuscular conditions involving TDP-43 proteinopathy.

## Materials and Methods

### Transgenic mice

A cDNA encoding mouse wild-type TDP-43 and EGFP linked by the internal ribosome entry site (*IRES*) was generated by standard PCR using the full-length mouse TDP-43 cDNA and EGFP as templates. The resulting cDNA (mTDP43-IRES-GFP) was cloned into the XhoI site of the MoPrp.Xho plasmid (ATCC#JHU-2) [43]. After the sequence verification and tests in cultured cells, the prp-mTDP43-IRES-EGFP construct was linearized by Not1 and injected into the pronuclei derived from FVB/NJ mice. The founder mice were screened by PCR using the primers complementary to the GFP and vector DNA sequence: forward TGCTGCTGCCCGACAACCA and reverse ATAACCCCTCCCCCAGCCTAGA. The positive founders were bred with wild type FVB/NJ mice, and the offspring were sacrificed and characterized for their expression of GFP in the CNS. Their brain and spinal cord were examined under a fluorescence microscope. The lines were terminated if 3 to 5 animals from the line showed no detectable GFP fluorescence. The lines surviving this screen were further analyzed for their GFP expression levels by immunoblot. Two lines, 19 and 42, were selected for a further screen for motor phenotypes. Both lines developed incompletely penetrant, late-onset motor dysfunction and paralysis. Line 19 was successfully converted to a homozygous line and further analyzed in detail. This line has been deposited at The Jackson Laboratory and will be available as Stock No. 031609.

### Behavioral analysis

Mice were housed in the University of Massachusetts Medical School animal facility managed by the Department of Animal Medicine. This facility is a specific pathogen-free (SPF) facility. Each cage housed 1-5 animals *ad libitum*. The rooms were maintained at 20-22°C and with a 12-12 light-dark cycle. All the behavioral experiments were approved by IACUC and conducted according to University of Massachusetts Medical School policies and procedures regulating the use of animals in research and the provisions of the PHS/NIH Guide for the Care and Use of Laboratory Animals.

#### Home cage observation

Mice were observed daily on weekdays for their general health and motor behavior, and their body weights monitored biweekly. The disease stages were assigned as follows: pre-symptomatic (pre-sym), slightly weak (swk), weak (wk), and paralysis (par). At the pre-sym stage (usually <10 months), the motor behavior was indistinguishable from the nTg mice. At the swk stage (at ∼10 months), the mice showed slight foot-dragging in their gait. At the wk stage (∼11-14 months), the foot-dragging became readily observable, and the movement became noticeably slowed. At the paralysis stage, two or more limbs became paralyzed, and the mouse was incapable of locomotion. The paralysis stage was the endpoint of the experiment, and the mouse was sacrificed for tissue harvesting.

#### HomeCageScan

To monitor the mouse behavior in their home cage continuously, we used the HomeCageScan system as previously described [44, 45]. Briefly, mice were housed individually in polycarbonate cages with minimal bedding (about 200 ml). A digital video camera was mounted on one side of the wall. Each mouse was recorded for 24h per week, with 12h daylight and 12h dim red light, and then returned to its cage with its littermates. Video data were analyzed by HomeCageScan software (Clever Systems, Reston, VA, USA) to quantify travel. Travel measures the overall motor/muscle functions by recording the distance traveled in meters.

#### Accelerating rotarod

Transgenic animals and age-matched controls were tested for time on accelerating rotarod from 12 to 72 RPM over three trials with a maximum time of 300s per trial at different age points. The longest time of three trials on the rotarod was recorded in seconds once the mice fell from the bar.

#### Grip strength

Transgenic animals and age-matched controls were tested for loss of four-limb grip strength using a grip strength meter at different time points. Mice were allowed to grip on a horizontal metal wire grid with four limbs. They were gently pulled back by their tails with steady force until they release their grip from the grid. Peak tension was recorded from five consecutive trials.

### Immunoblotting

Mice under deep anesthesia were decapitated. The spinal cord, brain, and other tissues were quickly harvested, snap-frozen in liquid nitrogen, and stored at -80°C. For protein preparation, frozen tissues were homogenized in a homogenization solution containing 25 mM phosphate pH 7.2, 1 mM EGTA, 1% SDS, 0.5% Triton X-100 and protease inhibitor mixture (Thermo Scientific) and heated at 95°C for 5 min. After clearing by centrifugation, protein concentration was measured using BCA assay (Pierce, Rockford, IL). The samples were heated in Laemmli buffer, and equal amounts of protein were loaded and resolved by SDS-PAGE. After transfer to nitrocellulose membranes, blots were blocked with 5% nonfat dry milk in PBST (0.25% Triton X-100 in PBS, pH 7.4) for 1 h, and then incubated with primary antibodies overnight at 4°C and then again with horseradish peroxidase–linked secondary antibodies (GE Healthcare) in PBST with 5% dry milk for 1 hour at RT. The dilutions and source of primary antibodies were as follows: GFP (Invitrogen G10362, 1:1000), rabbit polyclonal antibody raised to amino acids 394-414 of human TDP-43 (c-TDP43, custom made, 1:5000), α-Tubulin (Sigma,T5168, 1:10000), CNPase (Cell Signaling, #5664s,1:1000), MBP(Abcam,ab62631,1:1000), MCT1(Abcam, ab90582, 1:1000), NFkB-p65 (Cell signaling, #8242, 1:1000), and phosphorylated NFkB-p65 (Cell signaling, #3033, 1:1000). Membranes were washed three times, and proteins were visualized after ECL (Pierce) treatment and detected by the LAS-3000 imaging system (Fujifilm).

### Sedimentation assay

Mouse lumbar spinal cords were homogenized using a handheld polytron for 20 sec in lysis buffer (50 mM Tris-HCl, 150 mM NaCl, 0.5% deoxycholic acid, 1% Triton X-100, 20 mM NaF, 1 mM Na_3_VO_4_, 5 mM EDTA) with protease inhibitor (1:100 dilution, P8340, Sigma, St Louis, MO, USA) and phosphatase inhibitor cocktails (Thermo Fisher). The homogenates (100 µL/sample) were centrifuged at 12,000g at 4°C for 5 min. The supernatants were moved to new tubes and measured for protein concentration as described above. The pellets were rinsed 3 times with the lysis buffer and resuspended in 20 µL 1X Laemmli buffer. Ten micrograms of protein from the supernatant were mixed with 2X Laemmli buffer. The supernatant sample and an equivalent volume of pellet sample were heated at 95°C for 5 min, cleared by centrifugation, and then resolved by SDS-PAGE. The gel was then immunoblotted, as described above.

### RT-PCR and qRT-PCR

For total RNA extraction, frozen tissues or sorted cells were homogenized in cold TRIzol reagent (Invitrogen) following the manufacturer’s protocol. RNA was then reverse transcribed to cDNA using qScript cDNA SuperMix (Quanta BioSciences). For testing candidate splicing targets, RT-PCR amplification using between 33 and 37 cycles were performed from at least three nTg mice and three Tg mice. Products were separated on 2% agarose gels and visualized by staining with ethidium bromide and photographed. For qRT-PCR measurements of candidate gene targets, real-time PCR was performed on the cDNA using the primers for the targets. The PCR cycles were carried out in a Bio-Rad Real-Time PCR system (C1000 Thermal Cycler, Biorad), and the PCR product was detected using Sybr Green. The levels of target genes were standardized to the housekeeping gene GAPDH in individual animals and then further normalized to the mean ΔCT of the wild type mice.

### Immunofluorescence and immunohistochemistry

Mice under deep anesthesia were transcardially perfused with cold PBS, followed by 4% paraformaldehyde in PBS. The perfused mice were then immersed in the same fixative at 4°C for another 24-48h. After fixation, tissues were immersed in PBS containing 30% sucrose at 4°C for 2–3 days. Tissues were then frozen in OCT freezing media (Sakura, Torrance, CA) and stored at -20°C. Frozen sections were cut at 20 µm using a cryostat. For immunostaining, sections were incubated in the blocking solution (5% normal serum in PBS, pH 7.4) for 1 hour at room temperature (RT) and then incubated with a primary antibody in the blocking solution overnight at 4°C. The dilutions and source of primary antibodies were as follows: NeuN (Millipore MAB377, 1:200), calbindin (Millipore AB1778, 1:500), GFAP (Abcam Ab7260, 1:1000), IBA1 (BioCare Medical CP290AB, 1:200), APC (EMD Bioscience OP80-100UG, 1:200), GFP (Invitrogen G10362, 1:333), ChAT (Millipore AB1044P, 1:200), rabbit polyclonal antibody raised to amino acids 394-414 of human TDP-43 (custom made), TDP-43 (Encor biotechnology MCA-3H8, 1:250), NF-L (Cell Signaling, #2837,1:100), CNPase (Cell Signaling, #5664s,1:100), MBP(Abcam,ab62631,1:100), NFkB-p65 (Cell Signaling,#8242, 1:100), activated caspase3 (R&D system, AF835,1:1000). Sections were then washed 3 times for 5 minutes each and incubated in the appropriate secondary antibody at room temperature for 90 minutes. For immunofluorescence, the sections were washed 3 times in PBS for 5 minutes each and mounted with Vectashield mounting medium containing 4,6-diamidino-2-phenylindole (DAPI, Vector Laboratories) and sealed with nail polish. Images of the brain and spinal cord sections were taken with a confocal microscope (Leica).

For quantification of TDP-43 signal intensity in the nucleus and cytoplasm, sections were double-stained for TDP-43 and cellular markers. After staining, the cells were visualized and photographed using confocal microscopy. The cells in the ventral horn of the spinal cord were measured for their fluorescence intensity using the Nikon NIS Elements software. For each cell, the average fluorescence intensity was calculated. Cells on at least 5 different sections from each of the three or more mice per genotype were measured.

For immunohistochemistry, sections were washed 3 times in PBS containing 0.25% Tween 20 and then stained following the manufacturer’s instructions for Vectastain ABC kit, Elite PK-6100 standard ImmPact tm DAB peroxidase Substrate kit SK-4105 (Vector Lab). The sections were then mounted on slides and dried overnight at 55°C. After soaking in Xylenen 2 times for 2 minutes each, the slides were sealed with Permount (Vector Lab).

### Detection of demyelination

To detection of demyelination, mice were fixed by transcardial perfusion using 4% paraformaldehyde and 2.5% glutaraldehyde in 0.1M sodium phosphate (pH 7.6). Tissues were further fixed by soaking in the same fixative at 4°C for 24 hours. Luxol fast blue staining was performed on 10-μm spinal cord or brain paraffin-embedded sections for demyelination.

### Visualization and quantification of neurodegeneration

Visualization and quantification of the cortical neurons were carried out as described previously {Aliaga, 2013 #1266}. Briefly, whole mouse brains were placed in 30% sucrose solution for 2 days, frozen, and sectioned sagittally at 50 µm thickness. Layer V pyramidal neurons were counted using every ninth section, with a total of nine sections per half brain. Sections were mounted onto gelatin coated slides and stained with CTIP2 or Cresyl violet. Stereological counting was performed using Stereo Investigator software (MBF Bioscience, Williston VT). Counting was performed within the motor area of cortical layer V. Only pyramidal neurons with a soma greater than 15 µm in diameter were included. A single experimenter who was blinded to the genotype performed all counts.

For visualization of ventral root axons, mice were fixed by transcardial perfusion using 4% paraformaldehyde and 2.5% glutaraldehyde in 0.1M cacodylate (pH 7.6). Tissues were further fixed by soaking in the same fixative at 4°C for 24 hours. L4 and L5 roots attached to dorsal root ganglia were dissected and postfixed with 2% osmium tetroxide in 0.1 M phosphate (pH 7.6), dehydrated in a graded ethanol series, and embedded in Epon-Araldite resin. One-micron sections were stained with toluidine blue and examined and photographed by light microscopy.

For quantification of ventral horn motor neurons, lumbar enlargement of the fixed spinal cords was sectioned on a cryostat at 20µm thickness. Every other section was collected until a total of ten sections were collected from each spinal cord. The sections were stained with goat ChAT antibody at 4°C overnight. A secondary donkey anti-goat biotinylated antibody and a Vectastain ABC and DAB peroxidase Substrate kit (Vector Lab) were used to reveal motor neurons. Images of the spinal cord sections were taken using a Nikon microscope, and motor neuron numbers in the ventral horn region were counted manually from each section.

For muscle histology, isopentane in a container was pre-chilled with liquid nitrogen until the isopentane started to solidify at the bottom of the container. A fresh specimen isolated from gastrocnemius muscle was placed on a cork disc with a drop of OCT, which kept the muscle in the desired orientation. The specimen was frozen by immersion into the isopentane for about 5 seconds and then stored at -20°C. The frozen tissue was sectioned using a cryostat and stained with Hematoxylin & Eosin (H&E).

### Electromyography

Mice were anesthetized by inhalation of isoflurane. Animals were placed immediately on a heating pad to maintain their core temperature at 37°C. Measurements were performed using a Cardinal Synergy electromyography (EMG) machine. A ground self-adhesive gelled surface electrode was placed over the tail. Potentials were recorded from several sites of the muscles of all four limbs with a concentric needle electrode (30G) using a gain of 50 µV/division and a bandpass filter with low and high cut-off frequency settings of 20 and 10,000 Hz, respectively. The entire recording process took 15-20 minutes per mouse, after which the mice were euthanized by isoflurane overdose or used for tissue collection.

### Visualization and quantification of neuromuscular junctions

Animals are euthanized via an overdose of isoflurane, then transcardially perfused with PBS for 2 minutes, followed by 4% paraformaldehyde for 5 minutes. Gastrocnemius muscles were dissected out and placed in 1.5% paraformaldehyde for 24 hours at 4°C. The muscles were then washed with PBS for 30min at 4°C and placed in 25% sucrose overnight at 4°C. Muscles were embedded in OCT medium, frozen rapidly, and stored at -80°C. Sections were cut at 35nm thickness using a Leica Cryostat, placed on Superfrost Plus slides, and stored at -80°C. Slides were allowed to defrost for 30 minutes before use. Slides were washed 3 times with PBS for 5 minutes and then 3 times with 4% Triton X-100 for 5 minutes. Sections were blocked using 10% donkey serum in PBS for 3.5 hours at room temperature. A primary antibody solution of Rabbit anti-synaptophysin (ThermoFisher, 1:1000) and rabbit anti-Neuronal class III Beta-Tubulin (Biolegend, 1:1000) diluted in blocking solution was applied for 24 hours at 4°C. Slides were washed again with PBS. A secondary antibody solution of Alexa-488nm-labelled Donkey anti-Rabbit (ThermoFisher, 1:500) and Alexa-555nm α-Bungarotoxin (ThermoFisher, 1:500) diluted in PBS was applied overnight at 4°C in the dark. Slides were imaged on a Nikon microscope. Neuromuscular junctions were then counted and quantified as innervated or denervated Based on nerve occupancy of the endplates. Those with>50% occupancy was counted as innervated and those with <50% were counted as denervated.

## Results

### Prion promoter drives TDP-43 expression primarily in the CNS and causes age-dependent, progressive weakness, and paralysis

To determine the effects of an elevated level of TDP-43, we constructed a transgene in which the mouse prion promoter (Prp) [43] drives the expression of wild type murine TDP-43 cDNA. We employed the authentic TDP-43 sequence to avoid any possible effects of a tag. GFP was co-expressed from an internal ribosomal entry site (IRES) to monitor the transgene expression (S1A Fig). The founder lines were screened for their expression of GFP in the CNS. Two lines (lines 19 and 42) were obtained. Both expressed similar levels of GFP and TDP-43 (S1B Fig). Western blots of tissues from various organs showed that the transgene was predominantly expressed in the CNS (S1C, D Fig). Despite the readily detectable GFP expression, changes in TDP-43 levels were hardly noticeable in these lines compared with non-transgenic (nTg) mice (S1B Fig), possibly as a result of TDP-43 autoregulation, as previously reported [46]. A small cohort from each of the two lines was monitored for up to 750 days. Five of the ten mice from line 19 and four of the five mice from line 42 developed motor deficits ending in full paralysis (S1E, F Fig). The symptomatic mice also showed progressive weight loss towards the paralysis stage (S1G Fig), similar to other animal models for ALS [36].

To facilitate the analysis of these mice, we sought to elevate the gene dosage by making homozygous lines. We were unable to generate homozygous mice from line 42 but succeeded from line 19. As anticipated, the homozygous mice developed motor symptoms earlier than the original hemizygous line and became paralyzed before 400 days (Fig. 1A, B; S1 Video). This phenotype was completely penetrant. Males developed paralysis at younger ages (∼356±19 days) than the females (∼385±26 days) (Fig. 1B). The motor symptoms progressed over approximately three to five months before ending in paralysis based on several different behavioral measures, including body weight, travel velocity, grip strength, and rotarod performance (Fig. 1C-F, S1 Video). Given the accelerated and completely penetrant phenotype, we focused on these mice for further analysis and referred to the TDP-43 line 19 homozygous transgenic mice as TDP-43 transgenic (Tg) mice in the following text.

**Fig. 1.**
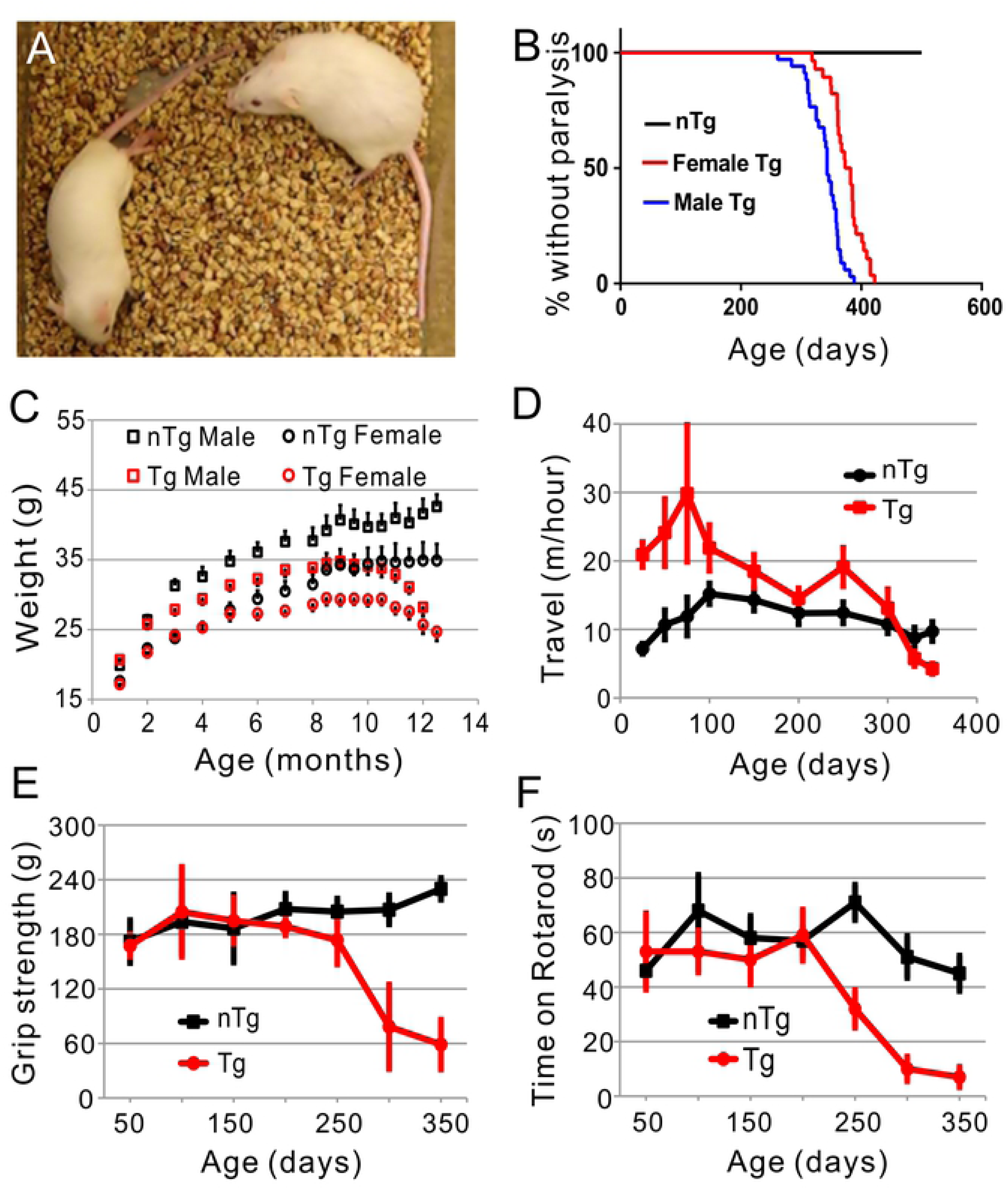
The TDP-43 homozygous transgenic (Tg) mice developed progressive motor phenotypes. (A) An example of Tg mice with hind limb paralysis (left). The mouse on the right was a non-transgenic (nTg) control. (B) Kaplan-Meier plot of Tg mice (37 males and 30 females) that are not paralyzed. (C) The bodyweight of the Tg mice peaked at ∼9 months of age and then declined until the paralysis stage. The values were averaged from 10 to 21 animals per group. (D) The home cage average travel velocity within a 24-hour period at various ages. The Tg mice went through a hyperactive stage before developing weakness and paralysis. (E) The Tg mice developed progressively weaker 4-limb grip strength after 250 days of age. (F) The Tg mice showed a declining rotarod performance after 200 days of age. Each data point in D, E, and F was an average of 16 animals.

As expected, the increased gene dosage in the homozygous mice increased the expression of the transgene compared with the hemizygous mice, as shown by the Western blots for GFP and TDP-43 (Fig. 2A). The hemizygous mice expressed TDP-43 at ∼10% above the nTg level in the spinal cord and ∼20% above in the frontal cortex (Fig. 2B). By comparison, the homozygous mice expressed ∼30% above the nTg level in the spinal cord and ∼40% above in the frontal cortex (Fig. 2B). Higher levels of mRNA were observed. The hemizygous mice expressed TDP-43 at ∼3 and ∼4 fold of the nTg levels in the spinal cord and the frontal cortex, respectively. By comparison, the homozygous mice expressed ∼5 and ∼8 fold of the nTg levels in the spinal cord and the frontal cortex, respectively (Fig. 2C).

**Fig. 2.**
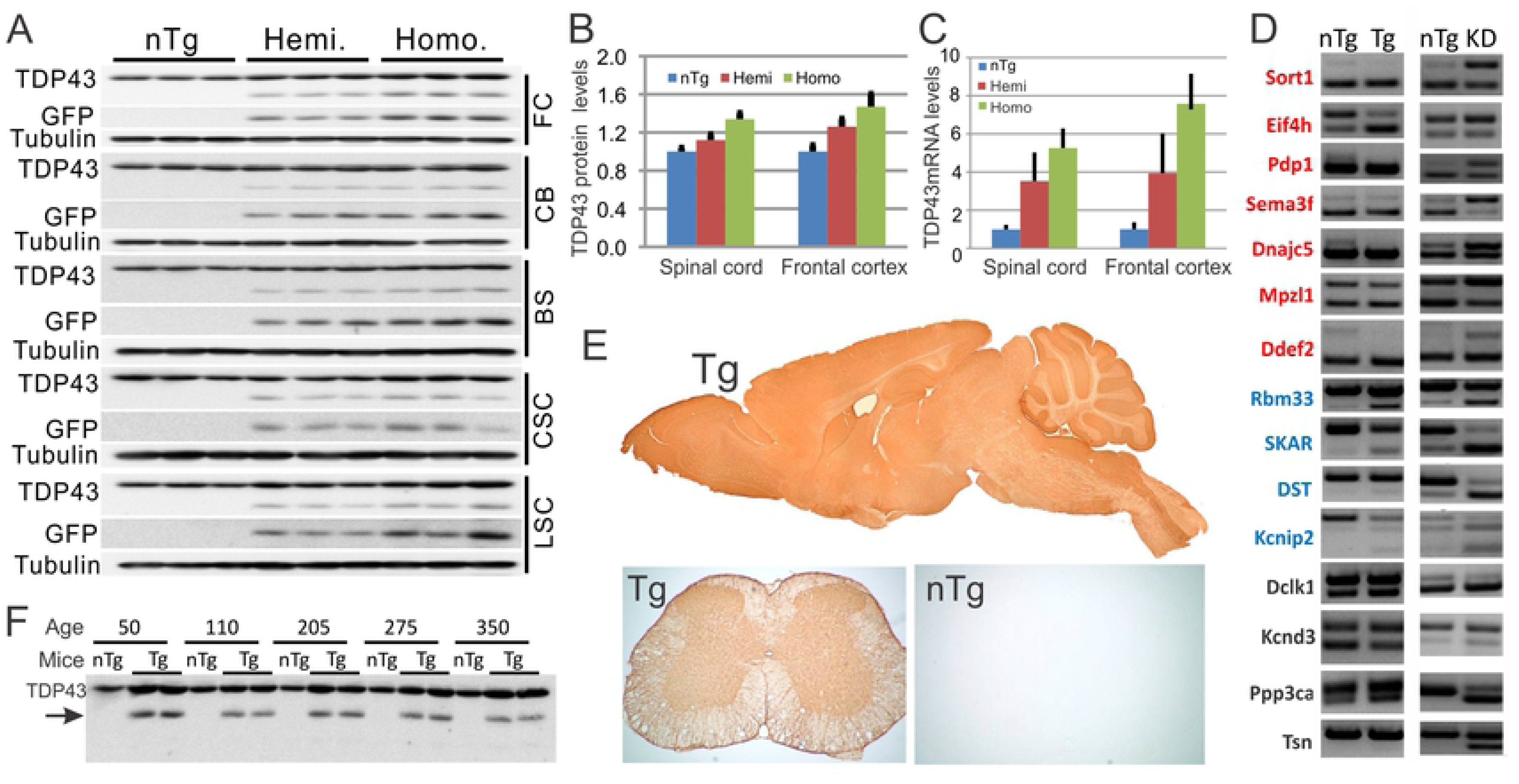
Transgene expression in the CNS of the hemizygous (Hemi.) and homozygous (Homo.) Tg mice. (A) A modest increase of TDP-43 protein in the frontal cortex (FC), cerebellum (CB), brainstem (BS), cervical (CSC), and lumbar spinal cords (LSC). Each lane represents one mouse. Notice that the 35-KD proteolytic fragment of TDP-43 was only detected in the transgenic mice but not in nTg controls. (B) Quantification of TDP-43 protein in (A) in FC and LSC. (C) Quantification of TDP-43 mRNA in FC and LSC. The number of animals used for FC quantification: nTg, 8; Hemi, 9; Homo, 11. For LSC: nTg, 9; Hemi, 12; Homo, 16. Error bars are standard deviation. (D) Comparing the alternative splicing patterns in the Tg mice vs. TDP-43 knockdown mice [45]. The PCR primers and the exon numbers are listed in S1 Table. (E) A sagittal section from a Tg mouse brain and a cross-section from either Tg or nTg mouse spinal cord were stained for GFP. Notice that the broad GFP staining throughout the Tg brain and spinal cord but not in the nTg. (F) Western blot showing stable TDP-43 protein levels in the TDP-43 Tg mice throughout different ages. The arrow point to the 35-KD fragment that is only present in the Tg mice.

TDP-43 is a gene-splicing modulator. To determine whether the modest elevation of TDP-43 level in the Tg mice impacted TDP-43 function, we measured the splicing patterns of some TDP-43-regulated mRNAs [47, 48]. For comparison, we also examined splicing patterns of the same target mRNAs in the TDP-43 knockdown (KD) mice [45]. Of the fifteen alternatively spliced genes that we measured, seven changed in the opposite direction in the TDP-43 Tg mice compared to the TDP-43 KD mice, with exon exclusion increased the Tg mice but decreased in the KD mice (Fig. 2D, red genes); four genes changed in the same direction, with exon exclusion enhanced in both the Tg and KD mice (Fig. 2D, blue genes); the final four genes did not change significantly in the Tg mice, but showed increased exon exclusion in the KD mice (Fig. 2D, black genes). Overall, the splicing modulation in our Tg mice was biased toward exon exclusion, consistent with the gain of TDP-43 function as reported in the literature [39, 47].

To determine where the transgene is expressed in the CNS, we stained brain and spinal cord sections for GFP. We observed a broad expression pattern in all regions of the CNS (Fig. 2E). The elevated level of expression was also sustained throughout the lifespan of the mice (Fig. 2F). To determine the cell types that express the transgene, we conducted double immunofluorescence staining. In spinal cord, we observed high levels of GFP in oligodendrocytes and astrocytes, and low levels in microglia and neurons, including motor neurons (Fig. 3). In brain, we observed relatively high levels of GFP in oligodendrocytes, astrocytes and neurons, and low levels in microglia (Fig. 3).

**Fig. 3.**
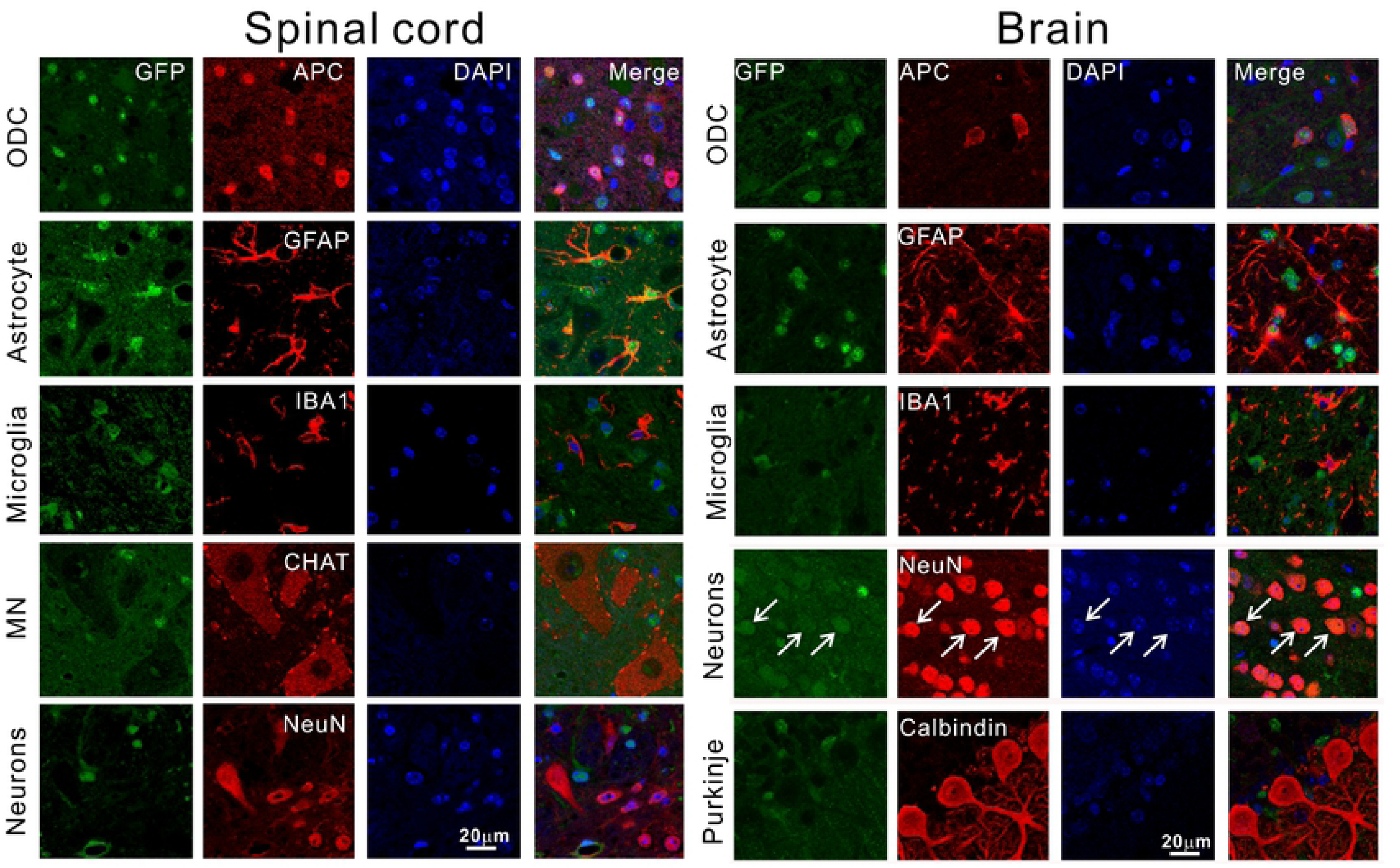
The Tg mice expressed transgenes in multiple different cell types in the spinal cord and the brain. In the spinal cord, the expression was relatively strong in oligodendrocytes (ODC) and astrocytes but weak in microglia and neurons, including motor neurons. In the frontal cortex (top four rows under the brain label), the transgenes are expressed strongly in ODC, astrocytes, and neurons (arrows) but weakly in microglia. Sections were double immunostained for GFP and various cellular marks as indicated. The transgenes were weakly expressed in Purkinje cells compared with the surrounding neuropils in the cerebellum. All panels are in the same magnification. All animals used are pre-symptomatic and less than 100 days old.

The increased TDP-43 expression may perturb the nuclear and cytoplasmic distribution of TDP-43. Because such perturbation may impact neurodegeneration [49], we quantified the nuclear and cytoplasmic TDP-43 staining intensities. TDP-43 was increased in both the nucleus and cytoplasm in all cell types that we measured (Fig. 4B). The overall increase was larger in glial cells than in neurons. In the nucleus, TDP-43 was up by ∼150-220% in glial cells, compared with up by ∼30-50% in neurons (Fig. 4B). Likewise, in the cytoplasm, TDP-43 was increased by ∼90-145% in glial cells, compared with an increase of ∼80-90% in neurons. Interestingly, the cytoplasmic increase was larger than the nuclear increase in neurons but was smaller in glial cells (Fig. 4B), resulting in a decreased cytoplasmic-to-nuclear TDP-43 in glial cells but an increased ratio in neurons compared with nTg mice (Table 1). Despite the increase, we did not observe TDP-43 aggregates in any disease stages, even though sedimentation experiments showed a ∼25% increase in the detergent-insoluble TDP-43 and ubiquitinated proteins in the Tg mice at the end disease stage (S2 Fig.), suggesting a modest increase in protein aggregation.

**Fig. 4.**
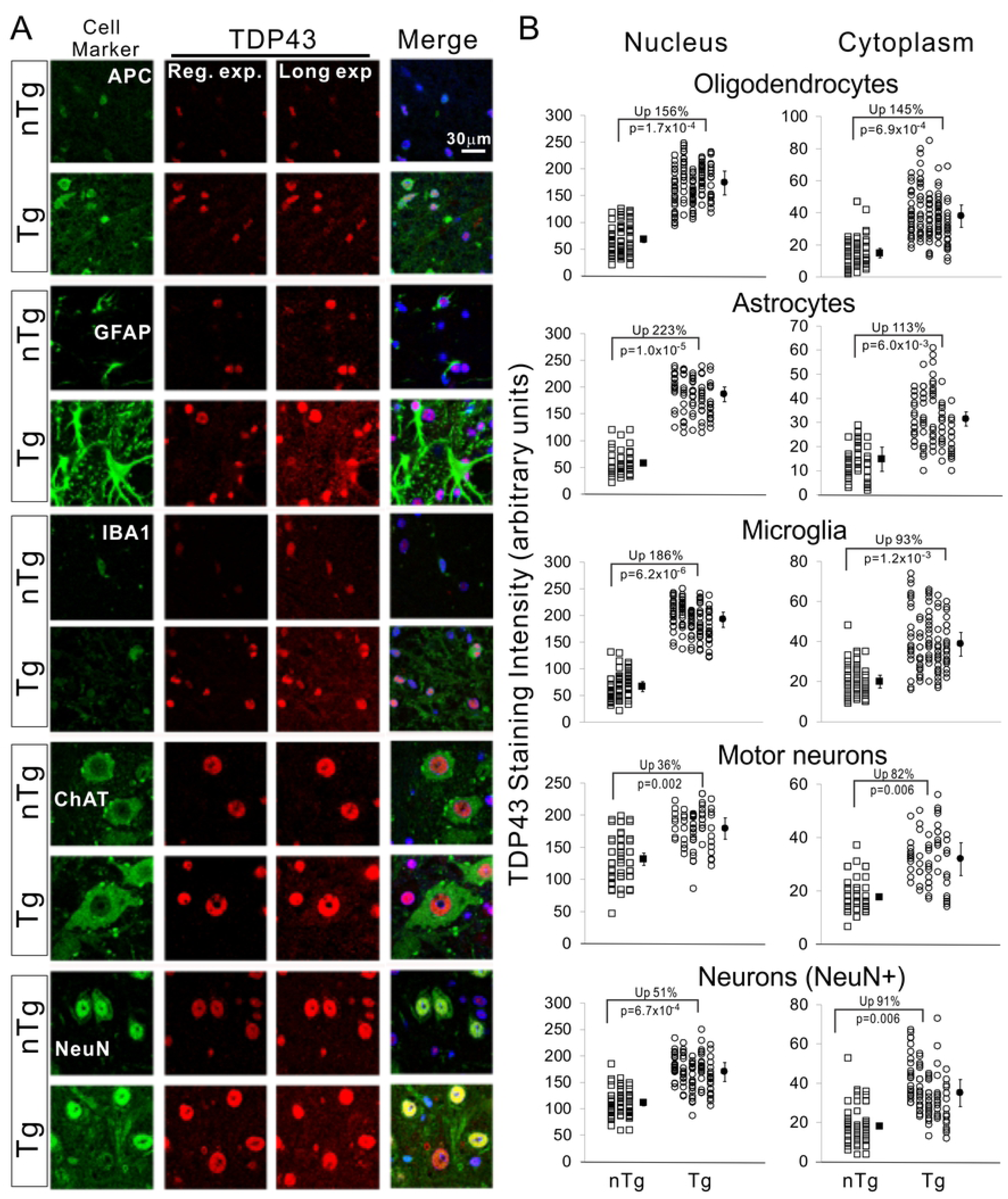
TDP-43 distribution in glial cells and neurons. (A) Mouse spinal cords were immuno-stained with cellular markers for oligodendrocytes (APC), astrocytes (GFAP), microglia (IBA1), motor neurons (ChAT), and pan neurons (NeuN). For TDP-43 staining, images of both regular (Reg.) and long-exposure (Long exp.) are shown. The long-exposure images were used to visualize the cytoplasmic signal and quantify TDP-43 staining intensity in the cytoplasm. All images are in the same magnification. (B) Quantification of staining intensity of TDP-43 in the nucleus and cytoplasm of various cell types show in (A). Each symbol represents measurement from one cell. Each column of symbols represents measurements from one mouse. The filled symbols represent averages for each genotype. The changes of averages in percentage in the Tg mice and the statistical p values are shown on the graphs. Student’s t test is used to obtain the p values. n = 3 for nTg and 5 for Tg mice, respectively.

**Table 1.**
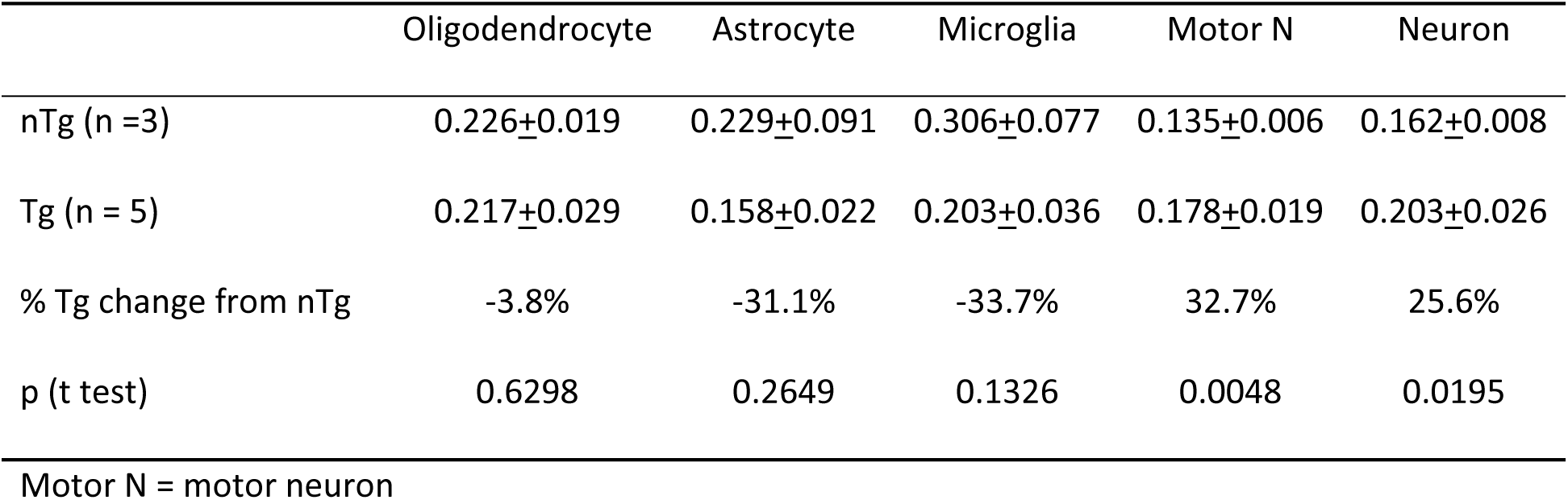
Ratio of TDP-43 cytoplasmic levels to the nuclear levels.

### TDP-43 mice develop severe white matter degeneration and oligodendrocyte injury in the low spinal cord

To determine the pathological basis of the clinical phenotype, we examined the spinal cord throughout the various disease progression stages in the Tg mice. The most conspicuous pathological feature was white matter degeneration. Luxol fast blue staining revealed normal myelination at age five months, but the staining became slightly pale at eight months. At ten months, the staining became paler (Fig. 5A). At the end-stage (12 months), the staining was completely lost in the ventral spinal cord (Fig. 5A, D and S3A Fig), indicating loss of myelin, which we confirmed by electron microscopy (S3B Fig). The myelin loss was accompanied by increasing astrogliosis (Fig. 5B, E) and microgliosis (Fig. 5C, F).

**Fig. 5.**
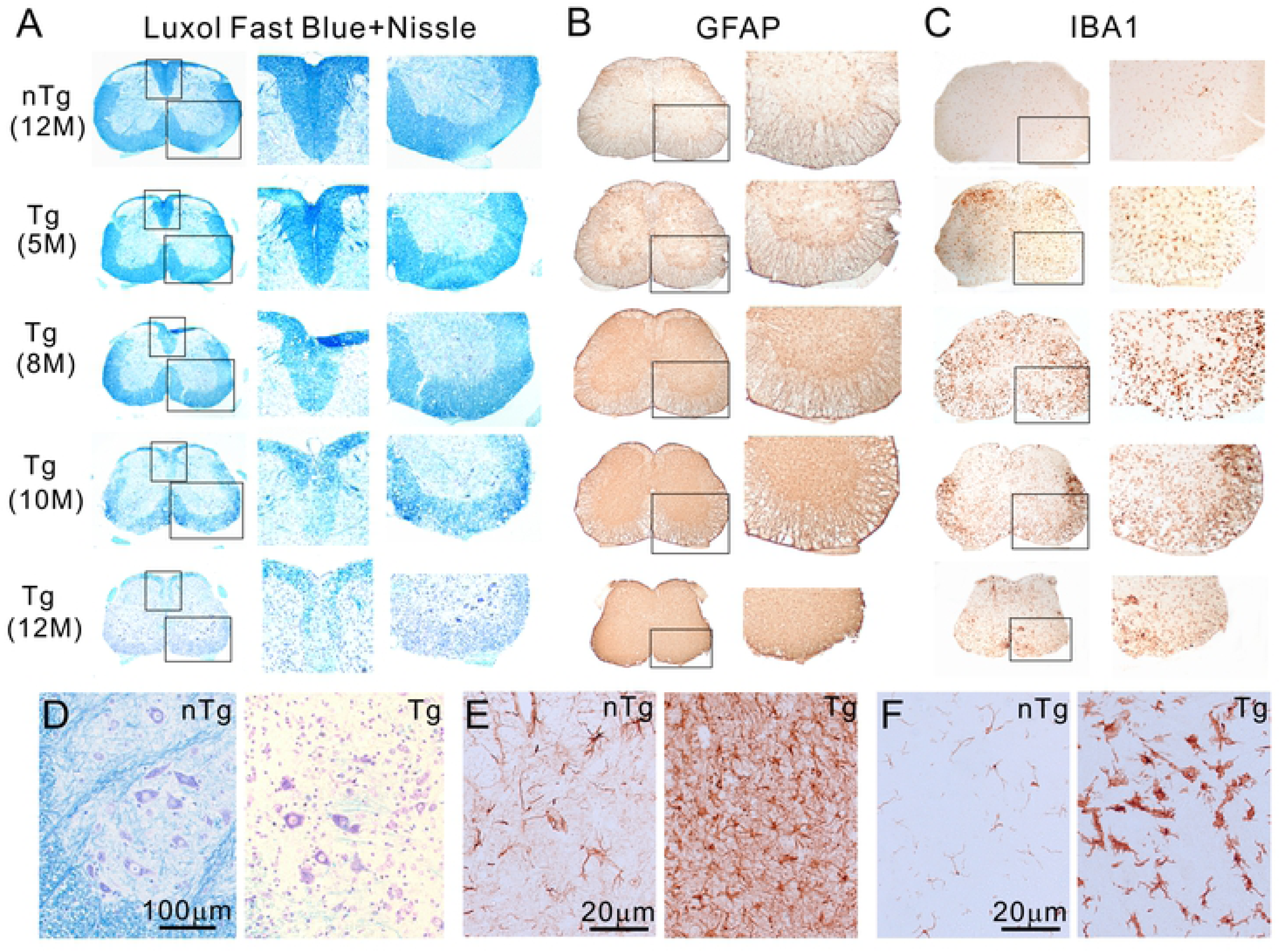
The Tg mice developed age-dependent myelin degeneration and gliosis in the spinal cord. (A) Late-onset, progressive demyelination, (B) astrogliosis and (C) microgliosis in the lumbar spinal cord of the Tg mice. The right panels are magnified views of boxed areas in the left panels. (D, E, F) Enlarged views of ventral horn stained with Luxol Fast Blue and Cresyl violet, GFAP and Iba1, respectively. Shown are images representative of five or more animals for each genotype and at each time point.

Myelin loss could be associated with injury in oligodendrocytes as a result of TDP-43 overexpression. Therefore, we doubly stained the spinal cord sections for oligodendrocyte marker APC and cell death marker activated caspase 3. We found an increasing association of activated caspase 3 and oligodendrocytes during the progression of the clinical phenotypes (Fig. 6A), and a reduction in oligodendrocyte-associated proteins, including CNPase, MBP, and MCT1 (Fig. 6B). Staining of CNPase in the white matter was also broadly lower than the nTg mice (Fig. 6C).

**Fig. 6.**
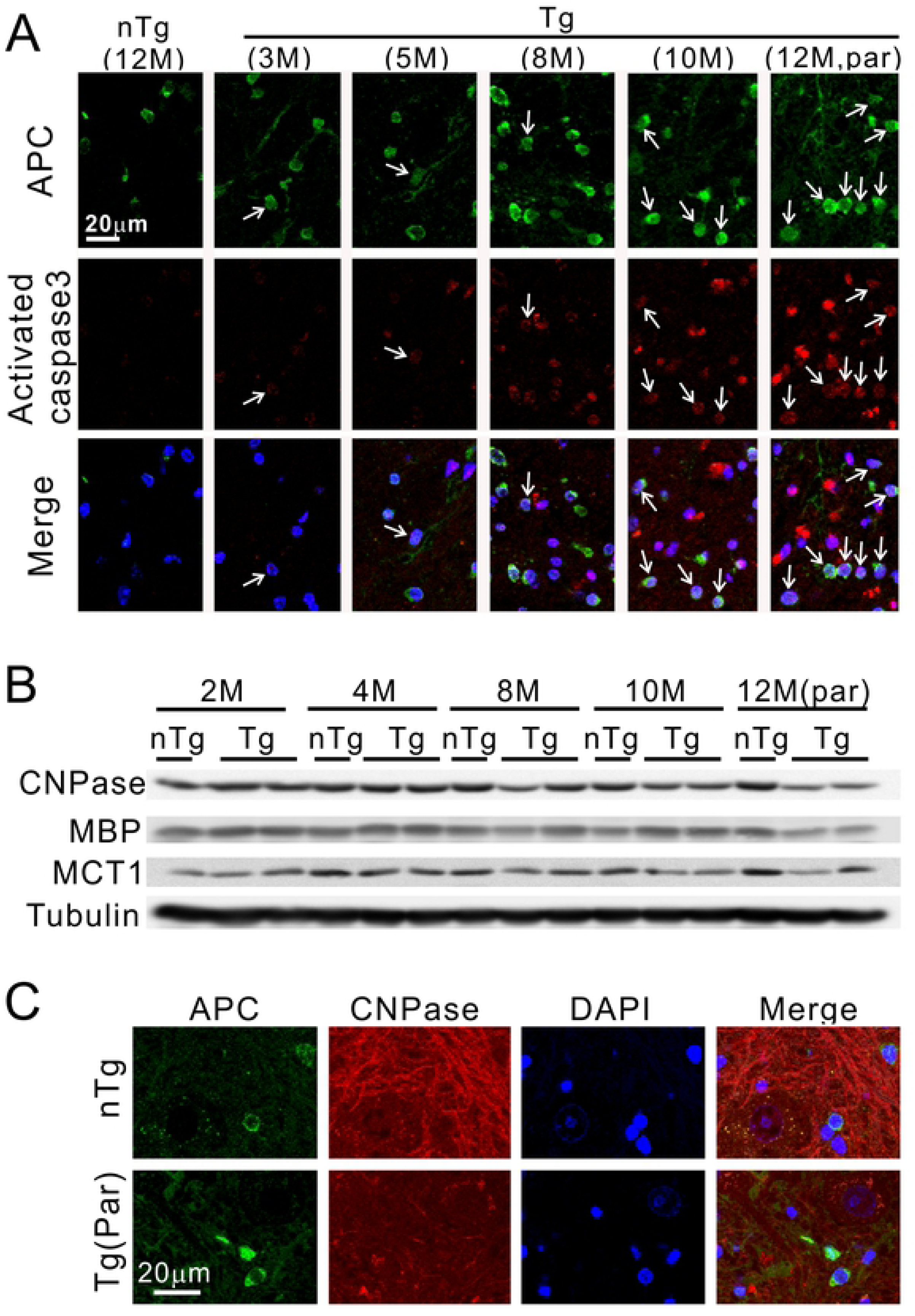
Oligodendrocyte pathology in TDP-43 mice. (A) Age-dependent and progressively increased activated caspase 3 signals are associated with oligodendrocytes (arrows) in the spinal cord of Tg mice. (B) Reduced myelin protein levels in the Tg mice with paralysis. M = month, CNPase = 2’,3’-cyclic-nucleotide 3’-phosphodiesterase, MBP = myelin basic protein, MCT1 = monocarboxylate transporter 1. (C) A reduced staining intensity of CNPase in the paralyzed Tg mice compared with the nTg mice.

### TDP-43 mice show no motor neuron loss but a mild muscle denervation

The progressive clinical motor phenotype suggests the presence of motor neuron injury. By Nissl staining and choline acetyltransferase (ChAT) staining, we did not observe gross morphological deviations from the controls (Figs. 5D, 7A). Quantification of ChAT-positive neurons in the ventral horn also did not reveal a significant difference between the Tg mice and the nTg mice (Fig. 7B). To verify this result, we examined and quantified ventral root axon numbers. We did not observe a significant change in the number of ventral root axons in the Tg mice from the nTg mice (Fig. 7C-E). Although we could not detect any motor neuron number changes, the Tg mice might have been at an early stage of motor neuron degeneration, which had not yet been reflected in the motor neuron numbers. To examine this possibility, we stained for activated caspase 3. We did not detect an association between the activated caspase 3 with motor neurons, although caspase staining of surrounding cells, possibly glial cells, were increased (Fig. 7F).

**Fig. 7.**
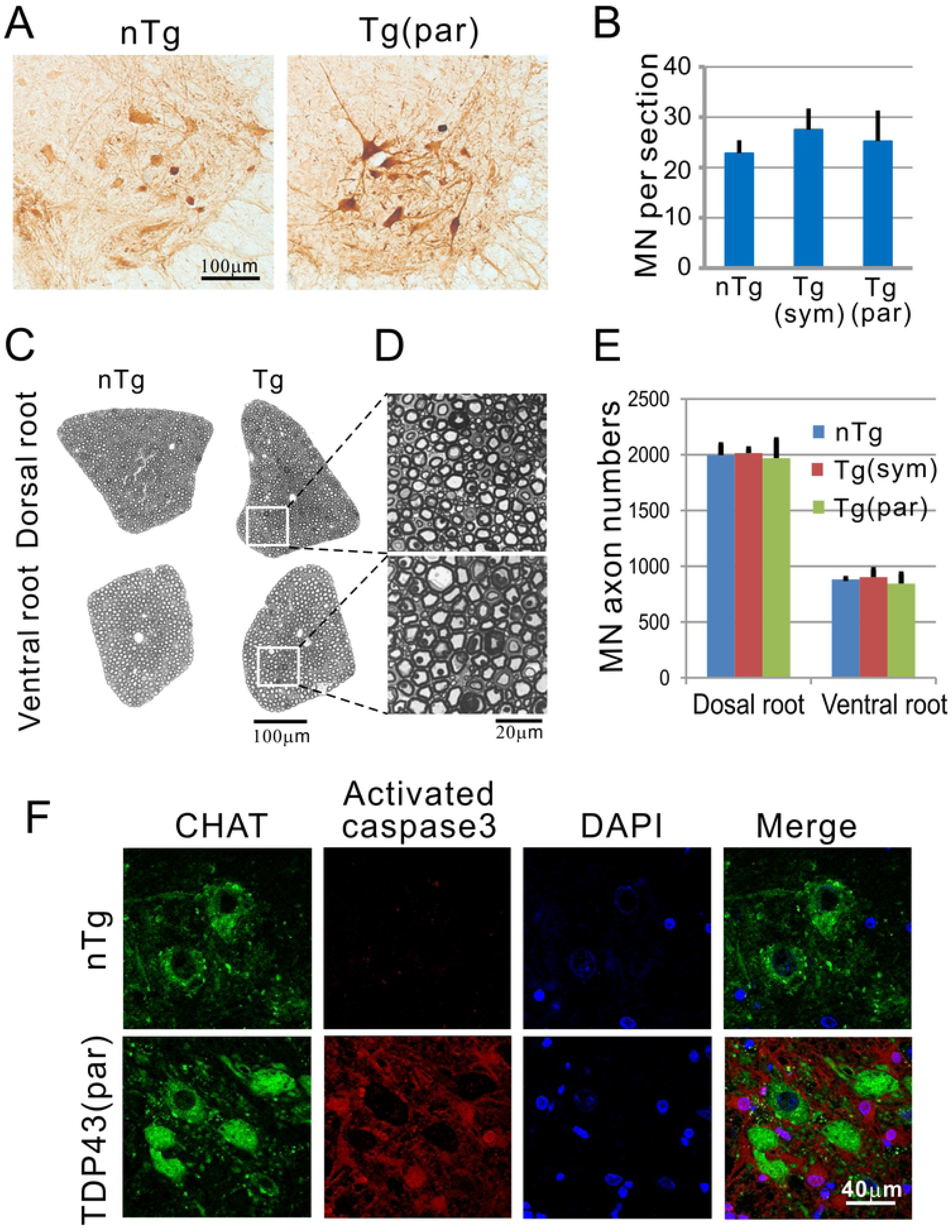
No spinal motor neuron loss in the ventral horn of the lumbar spinal cord. (A) ChAT staining of the ventral horn. (B) Quantification of ChAT-positive cells in the ventral horn of the spinal cord. ChAT-positive cells in the ventral horn were counted in 8-10 sections from the lumbar L4-5 levels of each animal. The average number per section was calculated from each animal. Four animals in each group were further averaged and shown here. Error bars represent standard deviation. (C) The axons in the ventral root and dorsal root remain normal. (D) High magnification of the boxed area in the dorsal and ventral root at the paralysis stage revealed no axon degeneration. (E) Quantification of dorsal and ventral roots showed no difference between Tg and nTg mice. (F) Immunofluorescence staining for activated caspase 3 show increased staining of the neuropils, but no motor neuron staining was detected. sym = symptomatic stage, par = paralyzed stage.

Despite the preservation of motor neuron cell bodies and their proximal axons, the distal axons and neuromuscular junction could be affected. To test this possibility, we examined muscle morphology and physiology. We did not observe muscle fiber atrophy (Fig. 8A, B). By needle electromyography (EMG), we observed that ∼half (3 of 5 homozygotes and 1 of 3 hemizygotes) of the paralyzed mice had a completely normal EMG pattern that was indistinguishable from the control nTg mice (Fig. 8C1). The other half of the paralyzed mice showed various degrees of positive sharp waves (PSWs) in different muscles ranging from normal (Fig. 8C1) to single and to multiple PSWs (Fig. 8C2, 3). Quantification of endplate nerve occupancy in the gastrocnemius muscle unveiled an average of ∼30% denervation in the paralyzed homozygous Tg animals compared with ∼10% denervation in the age-comparable control nTg mice (Fig. 8D).

**Fig. 8.**
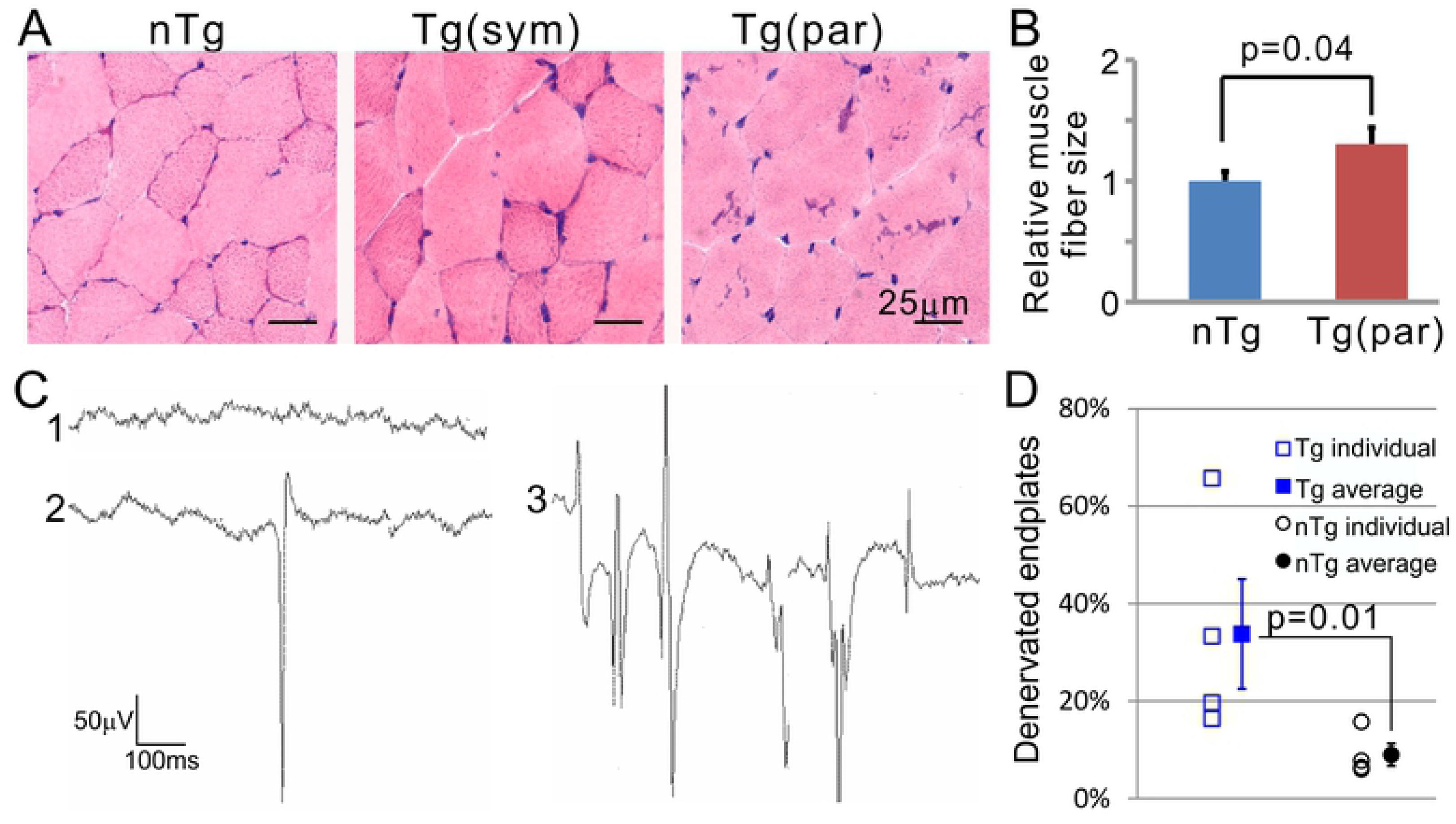
Denervation at the neuromuscular junction. (A) Examples of cross-sections of gastrocnemius muscle from 12-month old nTg (left), 10-month old (middle), and 12-month old paralyzed TDP-43 Tg mice (right). (B) Quantification of muscle fiber cross-section sizes. Muscle sizes from individual animals were averaged. The average values from four animals were further averaged and depicted. Error bars are standard errors. The p values were obtained by comparing the nTg with the Tg mice using student t test. (C) Examples of electromyographic (EMG) traces from an nTg mouse (trace 1) and a paralyzed Tg mouse (trace 2 and 3). (D) Quantification of denervated muscle endplates. n = 4 in both Tg and nTg groups. Statistical comparison between the Tg and nTg groups was conducted using Wilcoxon Non-parametric method.

### TDP-43 mice show elevated neuroinflammation in their spinal cord

Demyelination and gliosis are commonly associated with neuroinflammation [50]. Therefore, we examined the state of neuroinflammation in the TDP-43 transgenic mice. We found that the mRNAs of neuroinflammation-associated genes such as TNF-α and NF-κB p65 subunit were increased along with the disease progression (Fig. 9A). Furthermore, the p65 protein and its phosphorylation were also increased in parallel with clinical symptom development in the Tg mice compared with the nTg mice (Fig. 9B). By immunostaining, the increased p65 was associated with oligodendrocytes (Fig. 9C, arrows). These changes suggested the presence of neuroinflammation in the TDP-43 transgenic mice.

**Fig. 9.**
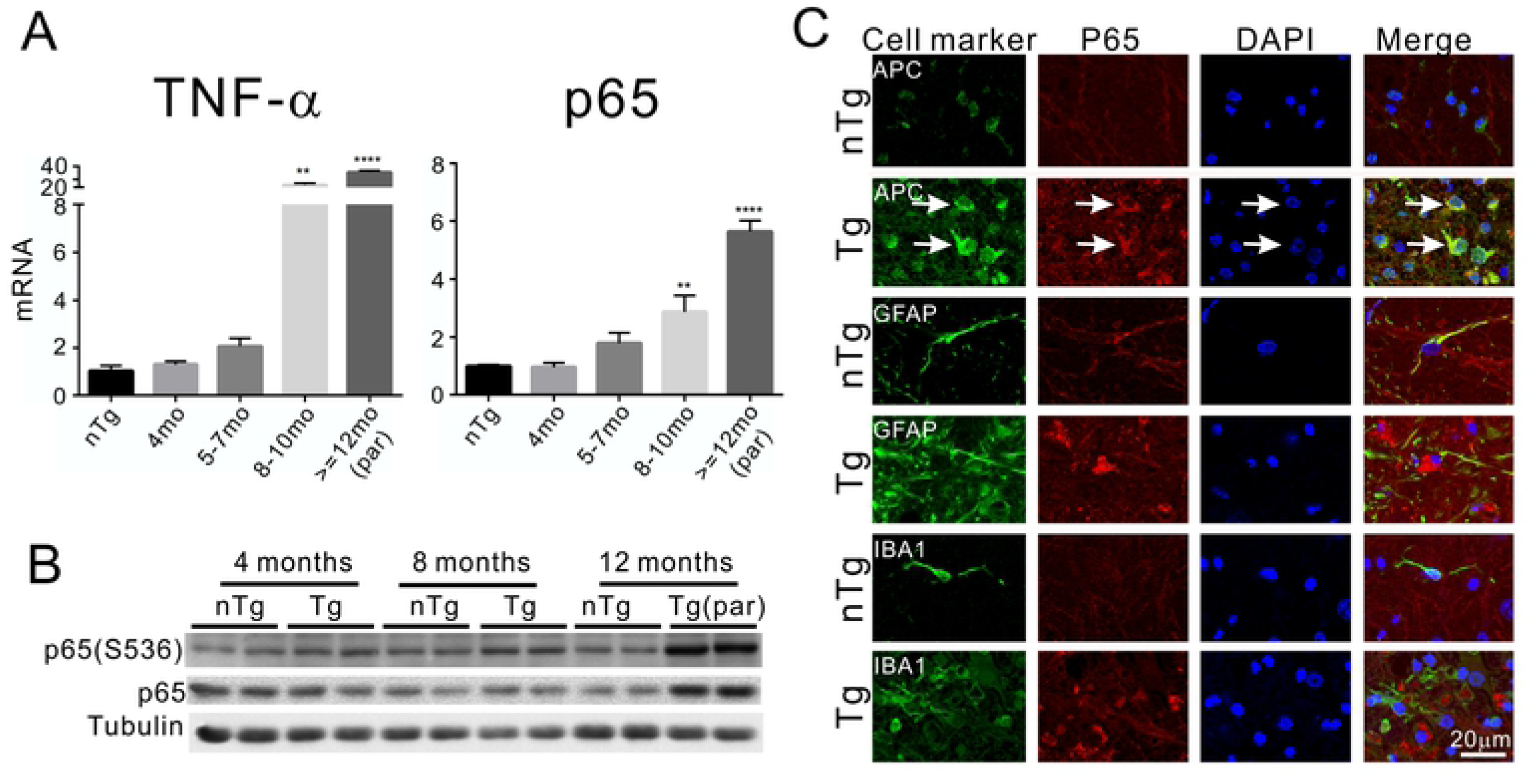
Activation of NF-kB pathway in the Tg mice. (A) TNF-α and p65 mRNA levels were dramatically elevated in the symptomatic Tg mice, and the peak was correlated with paralysis. The value of nTg mice is an average from eight animals with the age range from 102 days to 436 days. The nTg mice did not show an age-dependent change and therefore grouped together. Student t tests with Bonferroni correction were used to compare between the Tg and the nTg mice at different ages. n = 4-8 at each age for both groups. ** = p < 0.01, **** = p < 0.0001. (B) Western blot analysis showed both the phosphorylated p65 and the overall p65 levels were increased near symptomatic onset and paralysis stage in the Tg mice. (C) Immunostaining for p65 and markers of glial cells show localization of the increased p65 in the oligodendrocytes (arrows) but not in astrocytes and microglia in the Tg mice.

To further confirm this observation, we examined additional molecular markers for neuroinflammation. The expression of NADPH oxidase (NOX), which is involved in inflammation and oxidative stress, was dramatically increased. The three mRNAs of its subunits, gp91^phox^, p22^phox^, and p67^phox^, were increased by ∼600, ∼80, and ∼9 fold, respectively (Fig. 10A-C). Two genes involved in the production of prostaglandins, the inducible cyclooxygenase 2 (COX-2) and hemopoietic prostaglandin D synthase (HPDGS), were increased (Fig. 10D, E). GLT-1, an astrocytic glutamate transporter, was decreased at the end-stage (Fig. 10F). LCN2 and IL-6, two proinflammatory and neurotoxic cytokines, were also highly upregulated (Fig. 10G-I). Concomitant with these changes, the protein oxidation levels were increased (Fig. 10J).

**Fig. 10.**
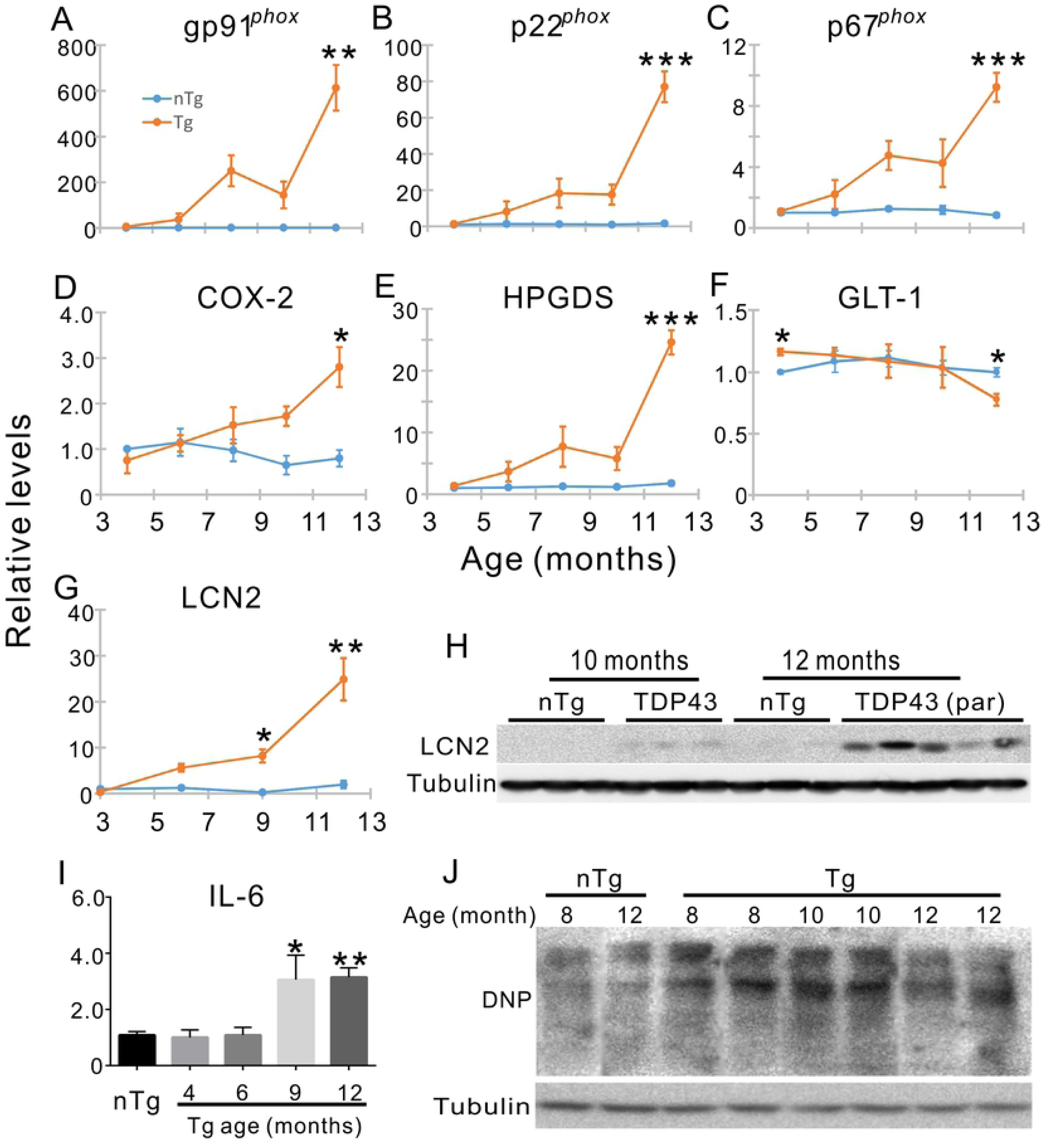
Age-dependent alterations in the expression of neuroinflammatory markers. (A-C) A dramatic increase in subunits of NADPH oxidase 2 (Nox2) gp91phox, p22, and p67 mRNA expression in late stages of motor dysfunction in the Tg mice. (E, F) Increased mRNA expression of prostaglandin-producing enzymes cyclooxygenase 2 (COX-2 or PTGS2) and hematopoietic prostaglandin D synthase (HPGDS), respectively. (G) Decreased expression of GLT-1 mRNA. (H, I) Increased expression of lipocalin 2 (LCN2) mRNA and protein, respectively. (J) Increased expression of IL-6 mRNA. (K) Western blot for dinitrophenyl (DNP) hydrazone derived from carbonylated proteins. Student t tests with Bonferroni correction were used to compare between the Tg and the nTg mice at different ages in (A-H) and (J). n = 3-8 at each age for both Tg and nTg groups. * p < 0.05, ** p < 0.01, *** p < 0.001.

Although not all the markers that we measured showed an increase (S4 Fig), the changes in the multiple neuroinflammation-associated markers suggest that the Tg mice undergo neuroinflammation in the spinal cord.

### TDP-43 mice show a mild white matter degeneration in CNS areas beyond the lumbar spinal cord

Because the TDP-43 transgenic mice expressed an elevated level of TDP-43 in the forebrain (Figs. 2A, E; 3), it is possible that the upper motor neuron degeneration could contribute to the paralysis phenotype. To examine this possibility, we investigated whether there was neurodegeneration in the forebrain. The brain weight in the TDP-43 mice did not differ from those of the nTg mice (S5 Fig), suggesting no widespread neuronal loss in the brain. Staining for myelin at different CNS levels showed that demyelination was most severe in the lumbar spinal cord and became progressively less severe towards the rostral CNS (Fig. 11A). The myelin staining was almost absent in the lumbar spinal cord, and paler compared to the nTg mice in the cervical spinal cord and lower brainstem (Fig. 11A). At the level of pons and above, no apparent difference in the staining was noticeable between the nTg and Tg mice (Fig. 11A).

**Fig. 11.**
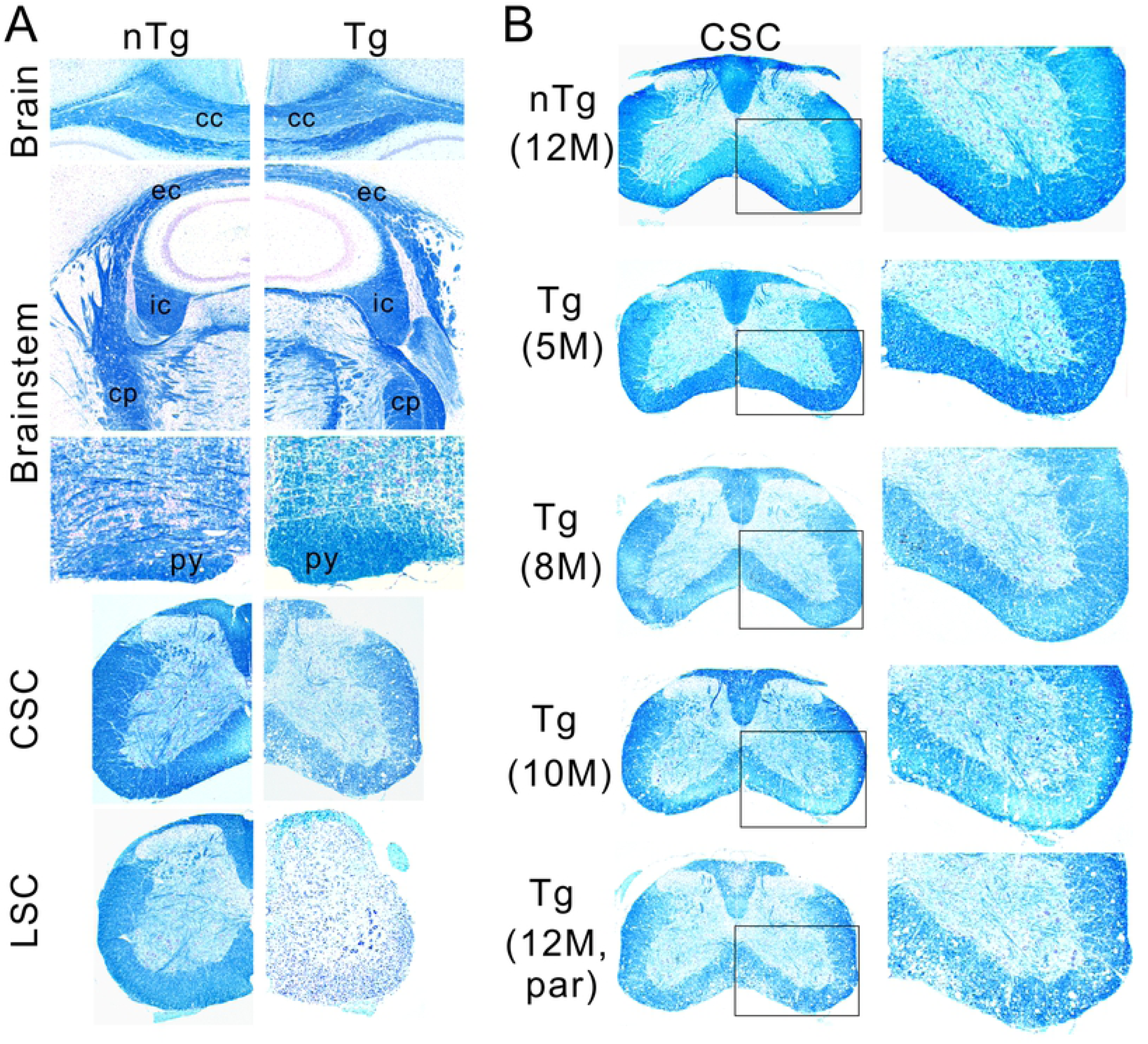
A survey of demyelination in the CNS. (A) The severity of myelin degeneration showed a caudal to rostral gradient in the CNS, with the lumbar spinal cord (LSC) being the most severely affected, followed by the cervical spinal cord (CSC), lower brainstem, upper brainstem, and corpus callosum (cc). py = pyramidal track, cp = cerebellar peduncle, ic = internal capsule and ec = external capsule. (B) The progression of demyelination in the cervical spinal cord. The panels on the right are enlargements of boxed areas in the panels on the left. Notice the degree of demyelination is less severe than in the lumbar spinal cord at 12 months old (compare with Fig. 5A and also between LSC and CSC in A of this figure). Shown are images representative of five or more animals for each genotype and at each age point.

At the cervical spinal cord level, the myelin staining intensity throughout the different ages showed a late-onset, progressive demyelination (Fig. 11B), a pattern similar to the lumbar region (Fig. 5A) though less severe. These results indicate that the overt demyelination in the Tg mice is mostly confined to below the brainstem.

We also surveyed gliosis in various regions of the brain in the paralyzed mice. We found elevated levels of astrogliosis and microgliosis in all regions, including the cerebral cortex, hippocampus, corpus callosum, internal capsule, cerebellum, and brainstem (Fig. 12), suggesting that elevated expression of TDP-43 provokes an inflammatory response by glial cells.

**Fig. 12.**
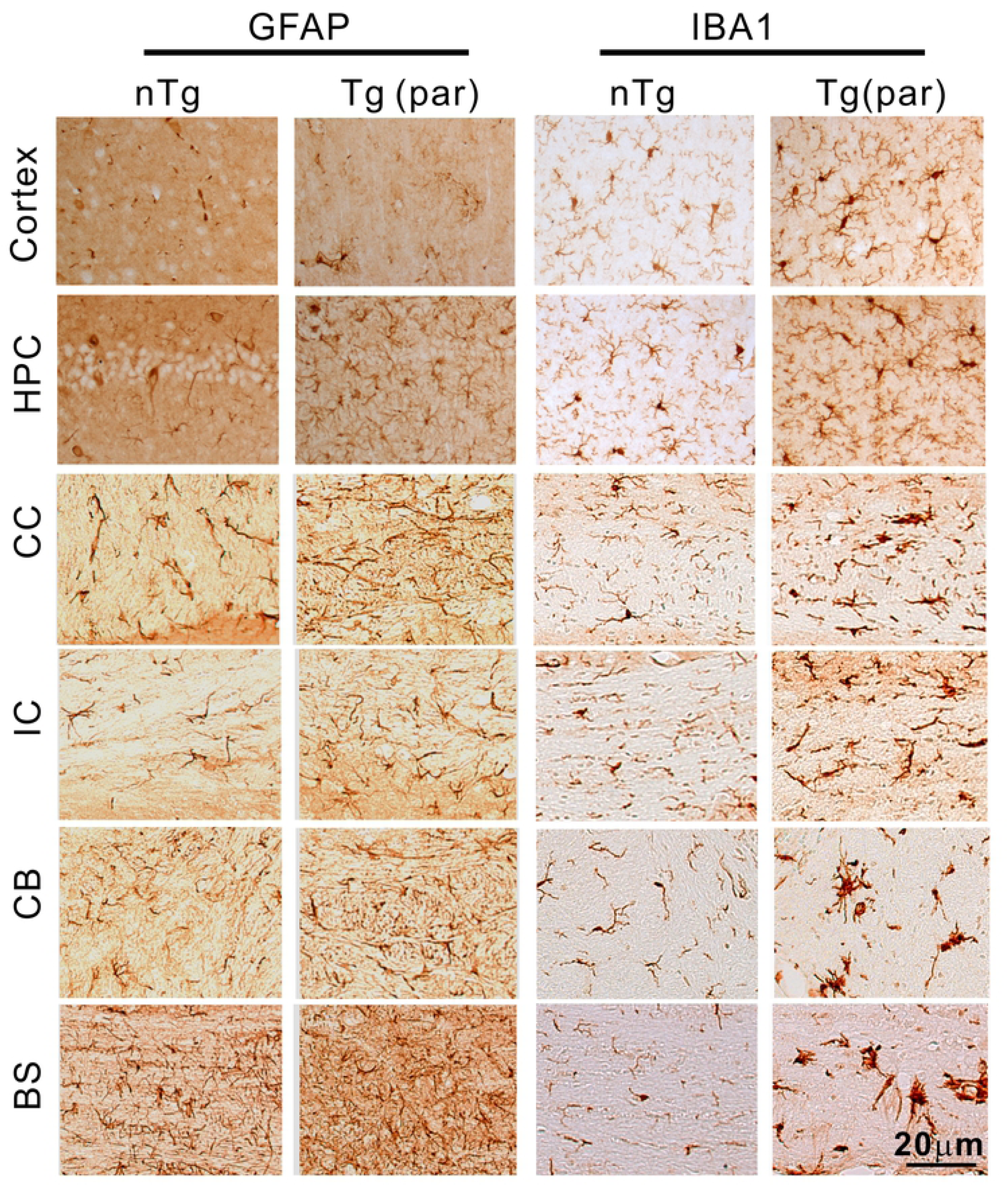
Mild but widespread gliosis in the upper brain regions in Tg mice. Staining for GFAP and Iba1 in the brain showed mildly elevated levels of astrogliosis and microgliosis in various regions of the CNS in the Tg mice, including the cerebral cortex, hippocampus (HPC), corpus callosum (CC), internal capsule (IC), cerebellum (CB) and brainstem (BS).

### TDP-43 mice develop neuronal loss in the motor cortex

Because TDP-43 was expressed in both neurons and glia in the cortex (Fig. 3), we investigated whether there was a neuronal loss in the motor cortex. Ctip2 staining revealed dramatically lowered staining intensity compared with the nTg control (S6A Fig). A quantification of Ctip2-positive neurons in layer V showed a near 50% loss (S6B Fig). However, because there was a significant loss of Ctip2 expression in the motor cortex of the paralyzed mice (S6C Fig), the loss of Ctip2 expression might have caused the lowered neuronal counts. Therefore, we applied Nissl staining (Fig. 13A, B) and repeated the quantification of neurons in layer V cortex. This approach revealed ∼30% neuronal loss (Fig. 13C). These results indicate that there is motor neuron loss in layer V of the motor cortex.

**Fig. 13.**
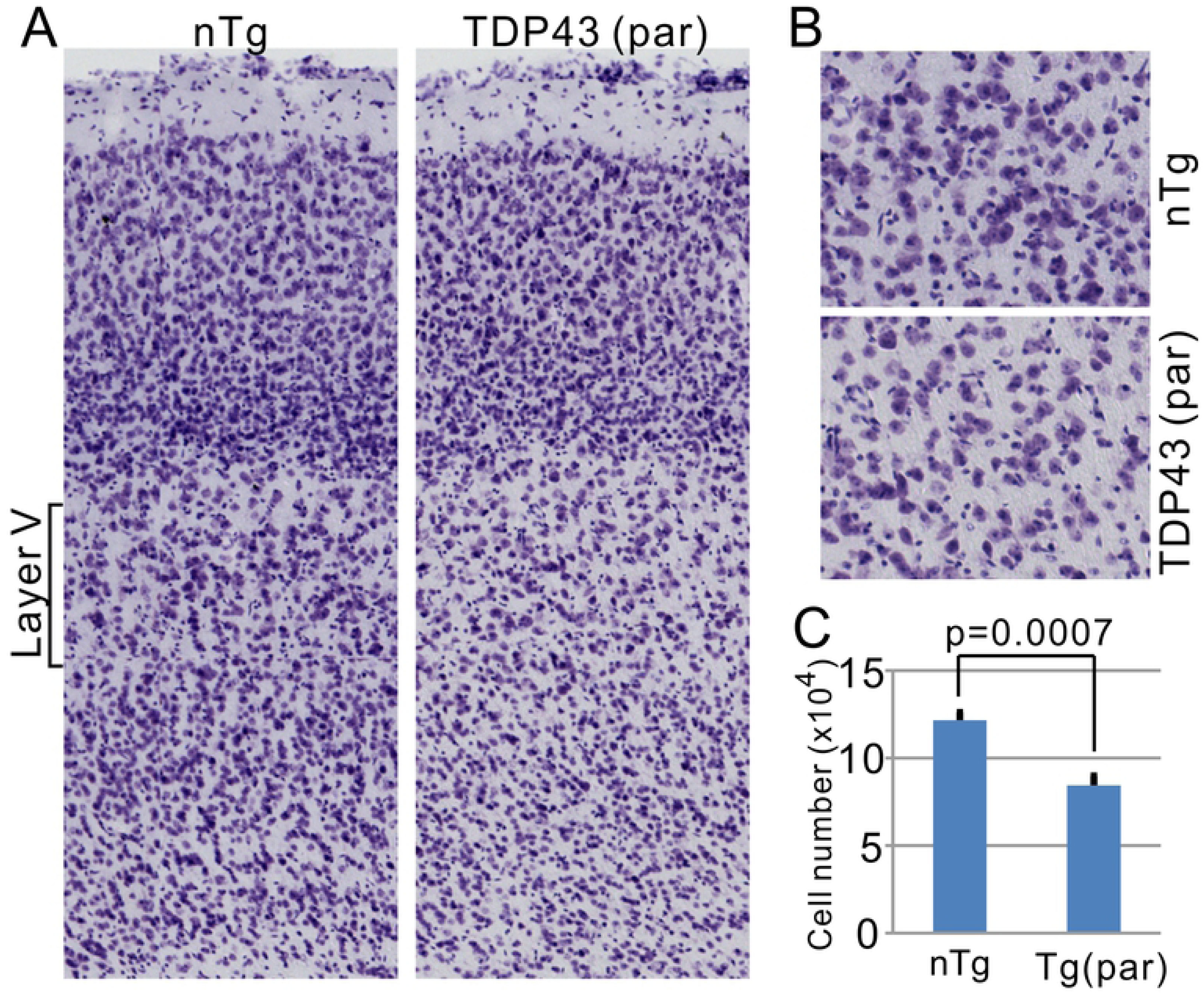
Visualization and quantification of Nissl-stained neurons in the motor cortex. (A) Nissl stained motor cortex in nTg and paralyzed Tg mice. (B) Enlarged views of layer V motor cortex. (C) quantification of large (>15 µm in diameter) Nissl-stained neurons in layer V. P value was derived from Student’s t test. n = 4 for nTg and 3 for Tg mice.

## Discussion

This study has established a line of TDP-43 transgenic mice that express exogenous TDP-43 protein at modestly elevated levels. Respectively for hemizygous and homozygous mice, the levels are ∼10% and ∼30% above the nTg level in the spinal cord and ∼20 and ∼40% increase frontal cortex (Fig. 2). These levels are comparable with the increase reported in human ALS and FTD [15, 28, 30, 34]. These TDP-43 mice show several key features of ALS, including late-onset progressive motor dysfunction ending in paralysis in mid-life, oligodendrocyte injury, demyelination, neuroinflammation, distal motor axon degeneration in the periphery, and cortical motor neuron loss. Thus, this transgenic mouse model demonstrates the multiple long-term adverse consequences of low-level TDP-43 overexpression *in vivo* and demonstrates that such low-level elevation of TDP-43, as observed in ALS and other neurodegenerative conditions, can trigger neurodegenerative clinical phenotypes.

How low-level overexpression of TDP-43 causes the late-onset neurotoxicity remains unknown. Pathological studies have consistently detected various forms of TDP-43 aggregates in ALS and FTD CNS tissues [51], suggesting an association between TDP-43 aggregation and the pathogenesis of these diseases. In vitro studies have shown that overexpression of TDP-43 can induce cytoplasmic liquid-liquid phase separation (LLPS), which, upon stress, transform into permanent aggregates. These aggregates can then draw TDP-43 out of the nucleus and kill the cell [52, 53]. However, in vivo models have shown divergent results. Some models show cytoplasmic TDP-43 aggregates either with or without nuclear TDP-43 depletion [37, 54, 55]. Other studies suggest that neither TDP-43 aggregation nor its nuclear depletion is required for neurodegeneration [39, 56]. Our results present a nuanced picture. Although we did not observe cytoplasmic aggregates and nuclear depletion in cells, we detected an increase in the detergent-insoluble TDP-43 and high molecular-weight ubiquitinated protein species (S2 Fig.), suggesting there is TDP-43 protein aggregation, albeit modest, in the TDP-43 mice. These results suggest that a modest elevation of TDP-43 levels can lead to TDP-43 protein aggregation, which could contribute to neurodegeneration in the TDP-43 mice.

Previous studies suggest that TDP-43 aggregation and nuclear depletion may lead to gain of toxicity as well as loss of function that ultimately cause cellular degeneration, including both neurons and oligodendrocytes [22, 23, 26, 57–61]. Analysis of mRNA splicing in mouse models has shown that a loss of TDP-43 function increases cryptic exon inclusion, whereas a gain of TDP-43 function leads to an increase in skiptic exons [9, 39]. Both the increases in the cryptic exon inclusion and the skiptic exons have been observed in human ALS samples, supporting the dual gain- and loss-of-function hypothesis [9, 39]. Consistent this hypothesis, both loss- and gain-of-function mouse models, including the model described in this report, produce motor neuron degeneration and ALS-like phenotypes [36, 39, 45, 62, 63], suggesting that each type of these models represent one aspect of the mechanism.

An intriguing observation in this study is that different cell types may be responding to TDP-43 overexpression differently. In oligodendrocytes, TDP-43 overexpression elicited similar percentage of increases in nucleus and cytoplasm (Fig. 4B), thereby maintaining the same cytoplasmic-to-nuclear TDP-43 ratio as in the nTg mice (Table 1). In astrocytes and microglia, the percentages of nuclear increase were approximately twice the cytoplasmic increases (Fig. 4B). As a result, the cytoplasmic-to-nuclear TDP-43 ratios were reduced, though not statistically significant (Table 1). By contrast, in motor neurons and non-motor neurons, the percentage of cytoplasmic increases was approximate twice the nuclear increases (Fig. 4B). Consequently, the ratio of cytoplasmic-to-nuclear TDP-43 was increased (Table 1). These observations suggest that neurons, particularly motor neurons, may handle the TDP-43 overexpression differently. It will be interesting to determine whether this observation can be replicated in other TDP-43 overexpression models in future experiments because of the potentially detrimental effects of an increased TDP-43 level in the cytoplasm [49].

Non-cell autonomous mutant toxicity is a well-established phenomenon in motor neuron degeneration provoked by mutant SOD1 [64]. Expression of mutant SOD1 in glial cells, including astrocytes, microglia and oligodendrocytes, accelerates motor neuron degeneration in vivo [65–69]. Glial cells expressing mutant SOD1 can also promote motor neuron degeneration in co-cultures in vitro [70–72]. Furthermore, mutant SOD1-expressing astrocytes and microglia secrete neuroinflammatory factors that are toxic and capable of killing motor neurons [66, 70, 71]. Mutant SOD1 expression in oligodendrocytes leads to cellular dysfunction, rendering them incapable of supporting neuronal axons [68, 69]. The evidence strongly supports the view that the glial expression of mutant SOD1 significantly contributes to motor neuron degeneration.

However, whether this is the case for TDP-43 is less clear. Astrocyte-specific expression of mutant TDP-43 in rats induces motor neuron degeneration as the mutant expression in motor neurons [73, 74]. However, mutant TDP-43-expressing astrocytes fail to show toxicity in motor neuron-astrocyte coculture or after being transplanted in rat spinal cord [75, 76]. Our TDP-43 transgenic mice showed two neurodegeneration patterns, one in the spinal cord and the other in the frontal cortex. In the spinal cord, the TDP-43 transgene was expressed highly in glial cells but lowly in neurons (Figs. 3, 4). The primary pathology was oligodendrocyte injury, demyelination, gliosis, and neuroinflammation (Figs. 5, 6, 9-11, and S3 Fig). However, no motor neuron loss was detected (Fig. 7). In the motor cortex, the TDP-43 transgene was expressed in both neurons and glial cells (Fig. 3). Approximately 30% of large pyramidal neurons were lost in layer V (Fig. 13 and S6 Fig). These results suggest that motor neurons can tolerate the detrimental effects invoked by modestly elevated levels of TDP-43 in their neighboring cells so long as the TDP-43 levels in themselves are maintained at relatively low levels. Even so, the tolerance of motor neurons to the injuries to their neighboring cells and neuroinflammation is probably limited, as illustrated by the presence of distal motor axon degeneration and neuromuscular denervation (Fig. 8). Because we have to sacrifice the mice at their paralysis stage, we do not know whether letting the disease progress further will eventually lead to the loss of the lower motor neurons. In any case, our results are consistent with contributions to motor neuron toxicity from both cell-autonomous and non-cell-autonomous sources.

A unique feature of our TDP-43 mice, compared with other established TDP-43 mouse lines, is the development of a fully penetrant progressive disease course (Fig. 1, S1 Video). The phenotype of late-onset, slowly progressing motor dysfunction to complete paralysis is reproducible for over >10 generations of homozygous breeding. This feature contrasts with other TDP-43 mouse models reported thus far. We note that there is a dose-dependent response to increased levels of TDP-43. In some reports, excessive expression of TDP-43 (e.g.,>3 fold of nTg level) causes acute toxicity and rapid demise of the animals. At the other end of the spectrum, models that express too little TDP-43 provoke no phenotype or at most mild phenotypes that require careful measurements to unveil [35–42]. Few transgenic lines developed motor dysfunction and paralysis but symptoms from the digestive system, highly variant phenotypes and lifespan (e.g., ∼1 to >15 months), and low penetrance complicate the phenotypes (e. g. ∼5% transgenic animals) [57, 77–79]. Thus, our TDP-43 mice provide a useful model for *in vivo* study of chronic TDP-43 toxicity derived from the modest elevation of TDP-43 levels and *in vivo* preclinical tests of experimental drugs targeting chronic TDP-43 toxicity in the CNS.

## Conclusions

We established a transgenic mouse model with a modestly elevated TDP-43 level. This model displays several ALS characteristics, including late-onset and progressive paralytic motor dysfunction ending in paralysis, neuroinflammation, and neurodegeneration. This study demonstrates that modest elevations in TDP-43 expression can trigger neurodegeneration and clinical phenotypes of ALS, suggesting that modestly elevated TDP-43 levels in humans could cause ALS and other neuromuscular disorders involving TDP-43 proteinopathy. Because of the easily observable, predictable, and progressive clinical paralytic phenotypes, this transgenic mouse model may be useful in preclinical trials of therapeutics targeting neurological disorders associated with elevated levels of TDP-43.

## Acknowledgments

The authors are grateful to the support from the core facilities at University of Massachusetts Medical School, including Transgenic Animal Modeling, Morphology, Electron Microscopy, Confocal Imaging, and Department of Animal Medicine. This work was supported by grants from the intramural research programs of National Institute on Aging (AG000946) to HC, by the National Natural Science Foundation of China, China (No. 81971200) to YG, by the ALS Association, the Angel Fund for ALS Research, ALS Finding A Cure, ALSOne, the deBourgknecht ALS Research Fund, and the Max Rosenfeld and Cellucci Funds for ALS Research, NINDS R01-NS088698 and RO1-NS111990 to RHB, and by NIH/NINDS RO1-NS101895, the ALS Association, and the Packard Center for ALS Research at Johns Hopkins to ZX.

## Supporting Information

**S1 Fig. Generation of transgenic mice that overexpress wild-type TDP-43.** (A) A cDNA encoding mouse TDP-43 and EGFP linked by internal ribosome entry site (*IRES*) were inserted into the backbone of MoPrp.Xho plasmid to generate the Prp-mouse TDP-43 transgene. The construct was composed of the following elements in linear succession: the Prp promoter, mouse wild-type TDP-43, IRES, EGFP gene, and poly A signal. This construct (Prp-TDP-43) expresses TDP-43 and GFP separately. (B) Western blot showed that the transgene was expressed in all the CNS regions in transgenic lines 19 and 42. FC, frontal cortex; CSC, cervical spinal cord; LSC, lumbar spinal cord; BS, brainstem; CB, cerebellum. (C) A survey of the transgene expression in different organs in transgenic line 19 showed that the transgenes were predominately expressed in CNS. Low levels of expression were also detected in heart (Hrt), lung (Lg) and kidney (Kdn). Other tissues are muscle (Msl), liver (Lv), and spleen (Spl). (D) A survey of the transgene expression in different organs in transgenic line 42. Similar to line 19, the transgenes were predominately expressed in the CNS. The samples in B, C, and D were prepared from animals between 55 and 65 days old. (E) A mouse at the paralysis stage. Its limbs were paralyzed, and the mouse lost its local motion capability. (F) Monitoring small cohorts of mice from lines 19 and 42 showed late-onset paralysis but incomplete penetrance from both lines up to 750 days. (G) Progressive weight loss in the aged mice of the two transgenic lines. The animal numbers are 3 to 4 for line 42, 3 to 5 for line 19, and 3 to 15 for non-transgenic (nTg) control mice at different age points. For the TDP-43 mice, only animals that develop paralysis were included in the weight plots. Error bars are standard errors.

**S2 Fig. Insoluble TDP-43 is increased in TDP-43 Tg mice.** (A) Western blot of detergent-soluble and insoluble TDP-43 and ubiquitinated proteins extracted from lumbar spinal cords of paralyzed Tg mice and age-matched nTg controls. Each lane was loaded with proteins from one animal. Arrows point to TDP-43 and its 35 KD and 25 KD fragments. Numbers on the right indicate molecular weights in kD. (B) Relative ratios of band intensity of the pellet over the supernatant. Bars represent averages of 7 nTg and 8 Tg animals in the TDP-43 quantification and of 4 animals in both nTg and Tg groups in the ubiquitin quantification. Student t test was used to compare between Tg and nTg mice. * indicates p<0.05 and *** p<0.001.

**S3 Fig. Widespread demyelination in the spinal cord of the Tg mice.** (Aa, e) cortical spinal track, (Ab, f) lateral funiculus, (Ac, g) ventral funiculus, (Ad, h) anterior commissure. (B) Electron microscopic images of ventral funiculus from a nTg mouse (left) and a Tg mouse (middle and right). Notice very few axons were wrapped by myelin in the Tg mice.

**S4 Fig. Inflammation markers that did not significantly change from the levels of nTg mice.** (A) cyclooxygenase 1 (COX-1 or PTGS1). (B) lipocalin type of prostaglandin synthase (LPGDS). (C) prostaglandin D2 receptor (DP1). (D) prostaglandin D2 receptor 2 (DP2). (E) inducible nitric oxide synthase (iNOS). (G) neuronal nitric oxide synthase (nNOS). Student t tests with Bonferroni correction were used to compare between the Tg and nTg mice at different ages. n = 3-8 at each age for both groups. No significance was found (p > 0.05).

**S5 Fig. No difference in the brain weight between the Tg and nTg mice** (n = 6 in both groups).

**S6 Fig. Visualization and quantification of CTIP2-positive neurons in the motor cortex.** (A) CTIP2 staining of the motor cortex in nTg and paralyzed Tg mice. The boxed areas in layer V are enlarged in the two panels on the right. Notice a substantial reduction in the CTIP2 staining intensity in the Tg mice. (B) quantification of CTIP2-positive cells in layer V. (C) Protein blot of CTIP2 in the motor cortex from nTg and Tg mice at different ages. P-value was derived from Student’s t test. n = 4 for nTg and 3 for Tg mice.

**S1 Table. PCR primers for data shown in** Figure 2D

**S1 Video. Tg mice developed progressive motor dysfunction and paralysis.** The transgenic mouse was marked with red ink on its tail. The other mouse in this video was a control nTg mouse. Notice the Tg mouse goes through stages from hyperactive to final paralysis.

